# Sex, age, tissue, and disease patterns of matrisome expression in GTEx transcriptome data

**DOI:** 10.1101/2021.03.09.434609

**Authors:** Tim O. Nieuwenhuis, Avi Z. Rosenberg, Matthew N. McCall, Marc K. Halushka

**Author notes:** Corresponding author at: Marc K. Halushka, M.D., Ph.D., Johns Hopkins University School of Medicine, Ross Bldg. Rm 632B, 720 Rutland Avenue, Baltimore, MD 21205. Email Addresses.

## Abstract

The extracellular matrix (ECM) has historically been explored through proteomic methods. Whether or not global transcriptomics can yield meaningful information on the human matrisome is unknown. Gene expression data from 17,382 samples across 52 tissues, were obtained from the Genotype-Tissue Expression (GTEx) project. Additional datasets were obtained from The Cancer Genome Atlas (TCGA) program and the Gene Expression Omnibus for comparisons. Gene expression levels generally recapitulated proteome-derived matrisome expression patterns. Further, matrisome gene expression properly clustered tissue types, with some matrisome genes including SERPIN family members having tissue-restricted expression patterns. Deeper analyses revealed 388 genes varied by age and 222 varied by sex in at least one tissue, with expression correlating with digitally imaged histologic tissue features. A comparison of TCGA tumor, TCGA adjacent normal and GTEx normal tissues demonstrated robustness of the GTEx samples as a generalized control, while also determining a common primary tumor matrisome. Additionally, GTEx tissues served as a useful non-diseased control in a separate study of idiopathic pulmonary fibrosis matrix changes. Altogether, these findings indicate that the transcriptome, in general, and GTEx in particular, has value in understanding the state of organ ECM.

## Introduction

The extracellular matrix (ECM) is the non-cellular scaffold found across tissues that provides structural integrity and mediates signaling to the cells it interacts with [1, 2]. The ECM is generated primarily by mesenchymal and other surrounding cells (fibroblasts, epithelial cells, adipocytes, etc.) during development and throughout the lifespan of an organism. It is primarily comprised of proteins, polysaccharides, and water, the exact type and proportions of each being unique to each tissue. This tissue specificity of the ECM is vital to the structure and function of each organ, with mutations of ECM genes resulting in Mendelian diseases that variably affects organs [3, 4].

While the study of ECM components predates cellular theory [5], the cataloging of its protein components has been a long endeavor. Due to the biochemical characteristics of ECM, such as its insolubility and frequent crosslinking, its composition has been difficult to interrogate even with the advancement of proteomic techniques [6]. The Hynes group catalogued a so-called “matrisome” using a combination of proteomic techniques applied to decellularized tissue samples and in silico bioinformatics analyses [7]. Their categorization is separated into two primary divisions, the core-matrisome proteins (n = 274) and matrisome-associated proteins (n = 753). The core-matrisome consists of three categories: collagens, proteoglycans, and ECM glycoproteins. These represent the structural elements of the ECM. The associated-matrisome, contrastingly, includes ECM regulators, ECM-affiliated proteins, and secreted factors.

While the majority of research in the ECM has been performed through mass spectrometry and immunohistochemistry, the field has not been comprehensively interrogated using transcriptomics. Herein, we generate a tissue-wide transcriptomic landscape of the core-matrisome and ECM regulators in normal human samples, and further investigate how the matrisome transcriptome is perturbed in lung disease and cancer. We use GTEx’s bulk sequencing data of 54 tissues comprised of 17,382 samples to create a generalized understanding of the similarities and differences of ECM expression between tissue types. GTEx’s phenotype data on the donor and sample level, including histological images, offers the opportunity to probe the effect of age and sex on matrisome expression. We also incorporate data from unrelated separate studies on cancer, available through The Cancer Genome Atlas (TCGA), and, in a third dataset, explore idiopathic pulmonary fibrosis (IPF). In summary, we demonstrated that GTEx gene expression data reasonably correlates with protein data and can serve as normal matrisome controls for disease states.

## Results

### Correlation of Matrix Gene and Protein Expression Patterns

Prior to utilizing the matrix transcriptome for analysis, we sought to establish the extent of correlation between the transcriptome and the more typically used proteome of the matrisome. For the purposes of this analysis we selected the core matrisome (glycoproteins, collagens, proteoglycans) and ECM regulators (n = 512 genes), and excluded the other matrisome-associated genes (ECM-affiliated proteins and secreted factors), which tend to not be structurally-related elements. Two large datasets of matched proteomic and genomic (bulk RNA-sequencing) expression data exist using either label free quantification (LFQ) [8] or a 10-plex tandem mass tag (TMT) approach [9] to quantify protein expression across multiple tissues. The LFQ dataset used intensity-based absolute quantification (iBAQ) derived from 29 human tissues, while the TMT was from 32 GTEx tissues. The LFQ proteomic approach had better correlations of protein expression to matched gene expression, highlighting challenges of normalizing TMT across samples (Fig. S1, Table 1), and therefore we focused on the LFQ dataset for further comparisons.

**Table 1.**
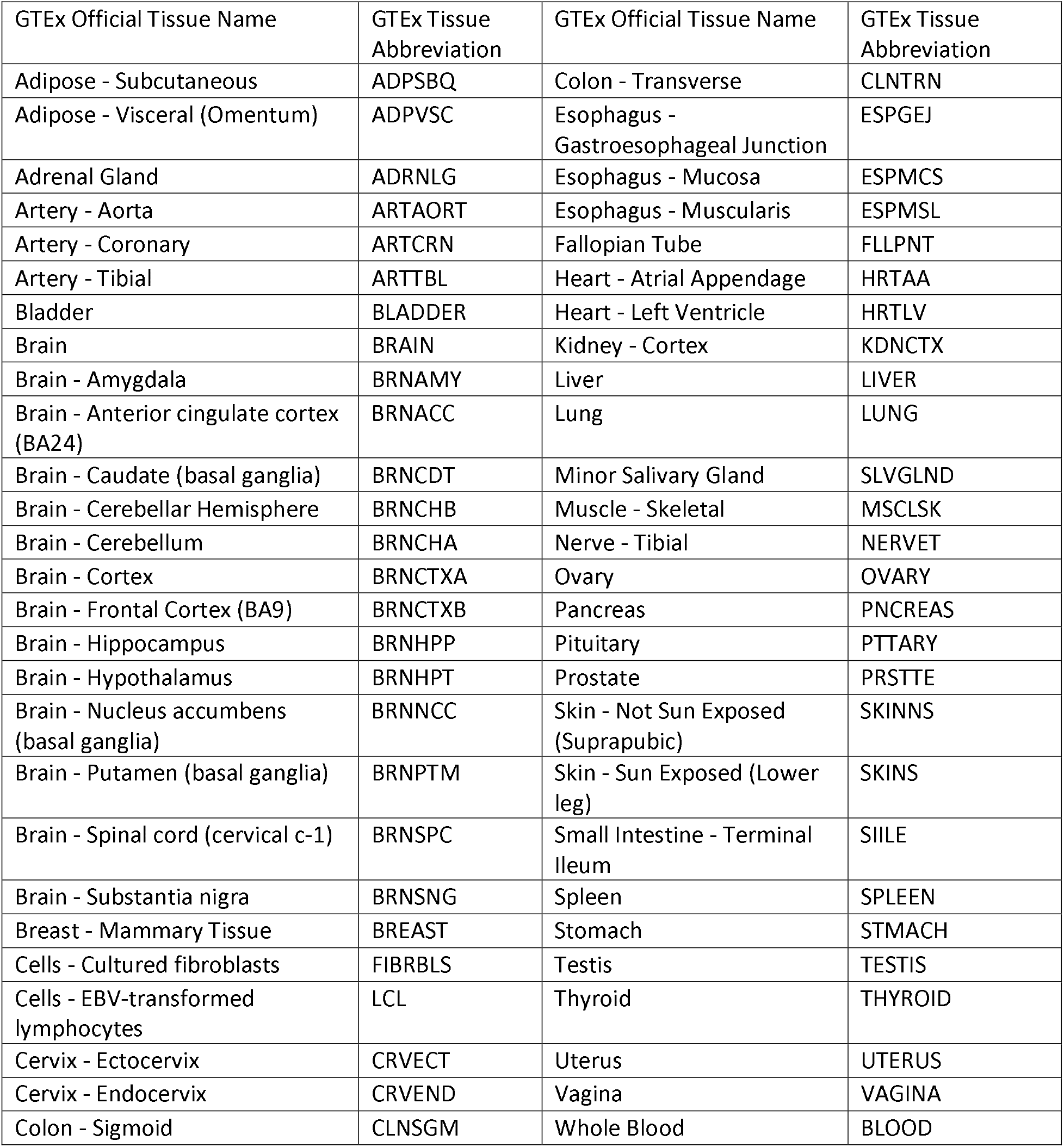
GTEx Tissue Abbreviation Dictionary.

**Table 2.** Lung sample clusters related to disease.

|  | Cluster 1 | Cluster 2 | Cluster 3 | Disease sum |
| --- | --- | --- | --- | --- |
| Normal Lung | 7 | 28 | 1 | 36 |
| Ventilator Injury | 8 | 2 | 0 | 10 |
| Acute Lung Injury | 3 | 5 | 0 | 8 |
| Idiopathic Pulmonary Fibrosis | 6 | 0 | 40 | 46 |
| Sample Sum | 25 | 35 | 41 |  |

For the 29 tissues, a matrisome correlation was generated using matrisome protein-gene pairs that had detectable levels of both the protein and gene. This ranged from 157 pairs in the pancreas to 237 pairs in the gallbladder (Fig. 1A). Using equal numbers of detected protein-gene pairs, 10,000 random sampled gene-protein pairs were performed to generate a Spearman’s ρ distribution (Fig. 1A). The matrisome Spearman’s ρ generally underperformed the permuted distribution (14/29 tissues had the matrisome correlation below 1 standard deviation (sd) from the mean of the distribution). The median average permuted correlation was 0.52, while the matrisome’s was 0.46. The matrisome Spearman’s ρ significantly correlated with the number of gene-protein pairs in each tissue (ρ= 0.773, p = 8.72e-07) indicating better correlation with increasing data depth. A deeper analysis of the matrisome classes demonstrate the overall lower performance was strongly the result of poor correlation of the ECM glycoprotein class (20/29 below 1 sd, 0/29 above 1 sd), which can be a challenging protein class to identify by general LFQ methods (Figure S2) [10]. Conversely and collectively, the proteoglycans, ECM regulators, and collagens had twice as many Spearman’s ρ above 1 sd (20) than below 1 sd (10).

**Fig. 1.**
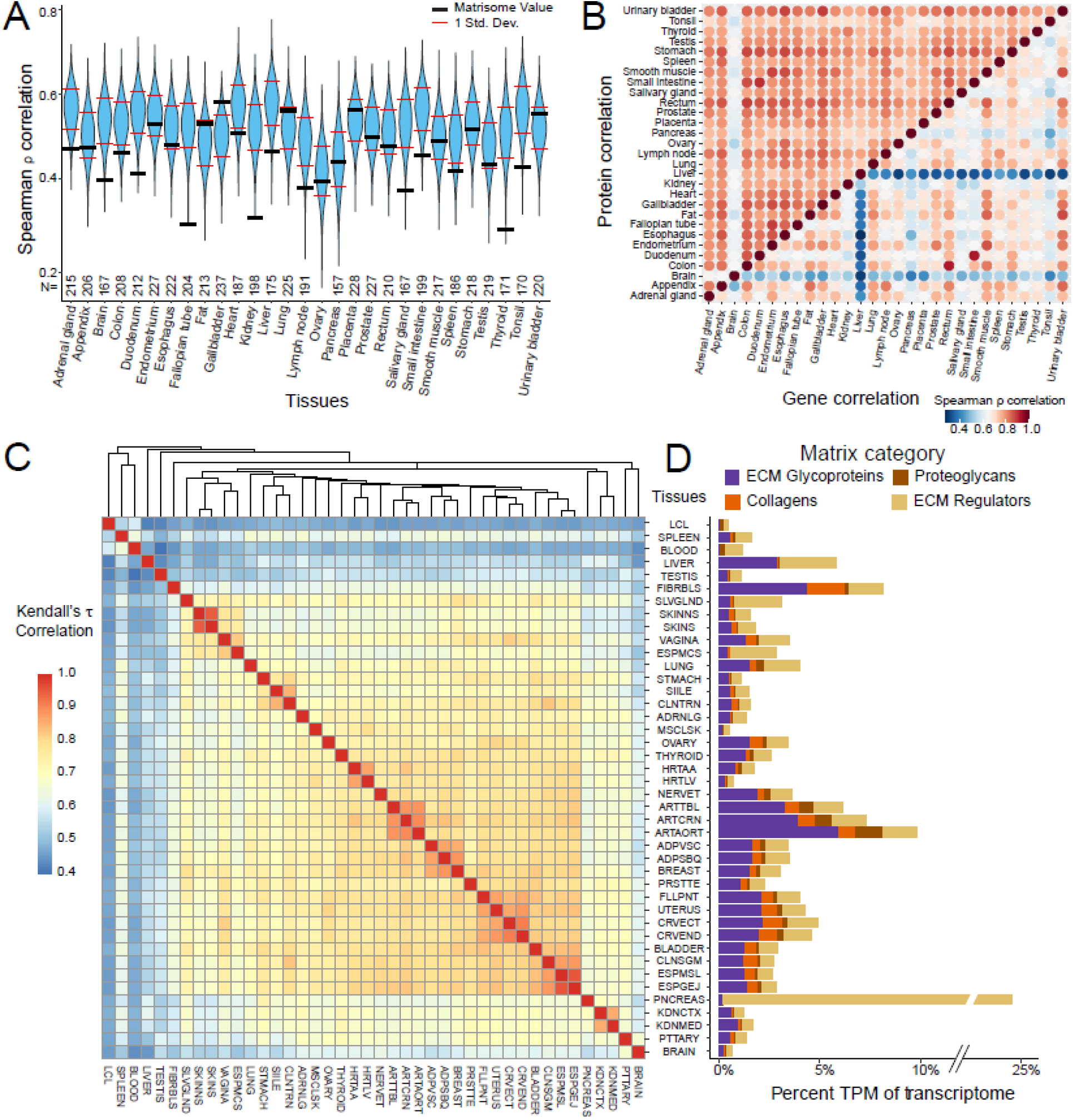
A transcriptomic and proteomic overview of the matrisome. A. A violin plot of 10,000 permutations of gene-protein Spearman’s correlations. The N of proteins per tissue is determined by how many protein/gene pairs were detected, with N being 203 for ovary and otherwise as reported. The black bars are the Spearman’s ρ correlation for the matrisome protein-gene pairs. B. A heatmap correlation of matrisome proteins (top left) and genes (bottom right) between tissues showing stronger correlations among proteins. C. A Kendall’s τ correlation heatmap of each tissues’ median expression of matrisome genes. Multiple brain region datasets are collapsed into one tissue by using the median expression of each gene across these samples. GTEx abbreviations are used. D. The median percent transcriptome of matrisome categories for each tissue. The colored bars each represent a different matrisome division.

From the original LFQ-based manuscript, intra-tissue correlations were much stronger in the complete proteomic dataset compared with the matched transcriptomic dataset [8]. When we selected matrisome components we observed strong similar protein intra-tissue correlations (Fig. 1B), potentially due to a short dynamic range of LFQ/iBAQ. Brain notably correlated least with other tissues. By contrast, there was significantly more diversity in correlation across the matrisome genes. Pairs such as rectum and colon or small intestine (jejunum/ileum) and duodenum were strongly associated (ρ = 0.931 and 0.934 respectively), as expected. More unrelated tissues such as the brain, pancreas, and liver had expected low correlation by matrisome gene expression. The pancreas separation is likely due to the high expression of regulatory matrisome genes in comparison to other tissue types [11], while liver has much higher levels of matrisomal glycoprotein expression [10]. These results demonstrated an overall reasonable correlation between matrix genes (transcriptomics) and proteins (proteomics) such that the evaluation of the GTEx transcriptomic dataset might prove valuable.

### The transcriptomic matrisome clusters GTEx tissue types

Using DESeq2 normalized bulk-seq GTEx gene expression data across 54 tissues and 17,382 samples, we generated an overview of the matrisome’s transcriptome. Using the median of normalized tissue counts, a Kendall’s τ correlation was generated between the 54 tissue types limited to the core and regulatory matrisome (n = 512 genes) (Fig. 1C, Table 1). There was strong clustering by similar organ types (skin, adipose, artery; Min τ = 0.95, 0.87, 0.89 respectively). Overall, cell type composition is a primary driver of additional tissue clustering based on the ratio of epithelium, smooth muscle, and other cell types [12].

We investigated genes that showed unique patterning between tissues and tissue types to better understand the inter-tissue expression that exists. Brain, for example, had enriched expression of proteoglycan stabilizers (*NCAN* and *BCAN*), while pancreas had highly-enriched expression of pancreatic enzymes *CELA3B, PRSS2*, and *PRSS3*. Organs containing stratified squamous epithelium, such as skin, esophagus, and vagina, shared higher expression of *SERPINA12, SERPINB13, SERPINB4*, and *LAMB4*, compared to other tissues. Glandular tissues, such as the small intestine and transverse colon also shared highly expressed matrisome genes including *MEP1A*, and *ADAMDEC1*. The liver notably had unique expression of genes including *ITIH2, SERPINC1, SERPINA10, F7, LPA* and *VTN*. The testis also had unique matrisome gene expression, although this may be the result of “leaky” transcription [13]. Of all sample types, the lowest global expression of matrisome genes was in EBV-transformed lymphocytes, while the liver and testis had the most uniquely expressed matrisome genes. To further understand this variation globally, we interrogated how the different matrisome sub-categories contributed to each tissue type.

We determined what percent of the transcriptome could be assigned separately to the divisions of matrisome-associated categories, ECM core and regulatory matrisome gene transcription (Fig. 1D). Overall, similar tissue types share the same level and composition of matrisome expression. For example, three adipose-dominant tissue types (ADPVSC, ADPSBQ, BREAST) had the same general composition and expression level of matrisome genes. The three arteries share similar compositions, with increasing matrisome transcript levels correlating with medial wall thickness [14]. EBV-transformed Lymphocytes (LCL) serve as a useful negative control, as they fail to generate matrix in culture. The pancreas remains an outlier due to its high expression of pancreatic enzymes that fall into the category of ECM regulators (*PRSS1, PRSS2, CELA3A*) [11].

### Some matrisome gene expression is modulated by age and associates with age-related histology

Aging is a biological process often linked to diseases such as cancer, heart disease, Type II diabetes, and osteoporosis [15, 16]. Aging impacts the matrisome through fibrosis and scarring, a component of which is collagen deposition. Using the limma package to regress gene expression on age while correcting for sex, we identified 1,667 instances of 388 unique core and/or regulatory matrisome genes associating with subject age in 40 tissues (Holm correction). Of these, 1,241 gene-tissue pairs increased with age while 426 decreased (Fig. 2A, Figure S3, Table S1). The most varied genes by age were *COL16A1, LTBP2, MGP* and *TNXB* found in 13-14 tissues each, although two of these genes did not have the highest expression change in any organ. The median number of tissues that genes varied in was 4. Of the 217 genes that had consistent directionality across tissues, 59 decreased and 158 increased with age.

**Fig 2.**
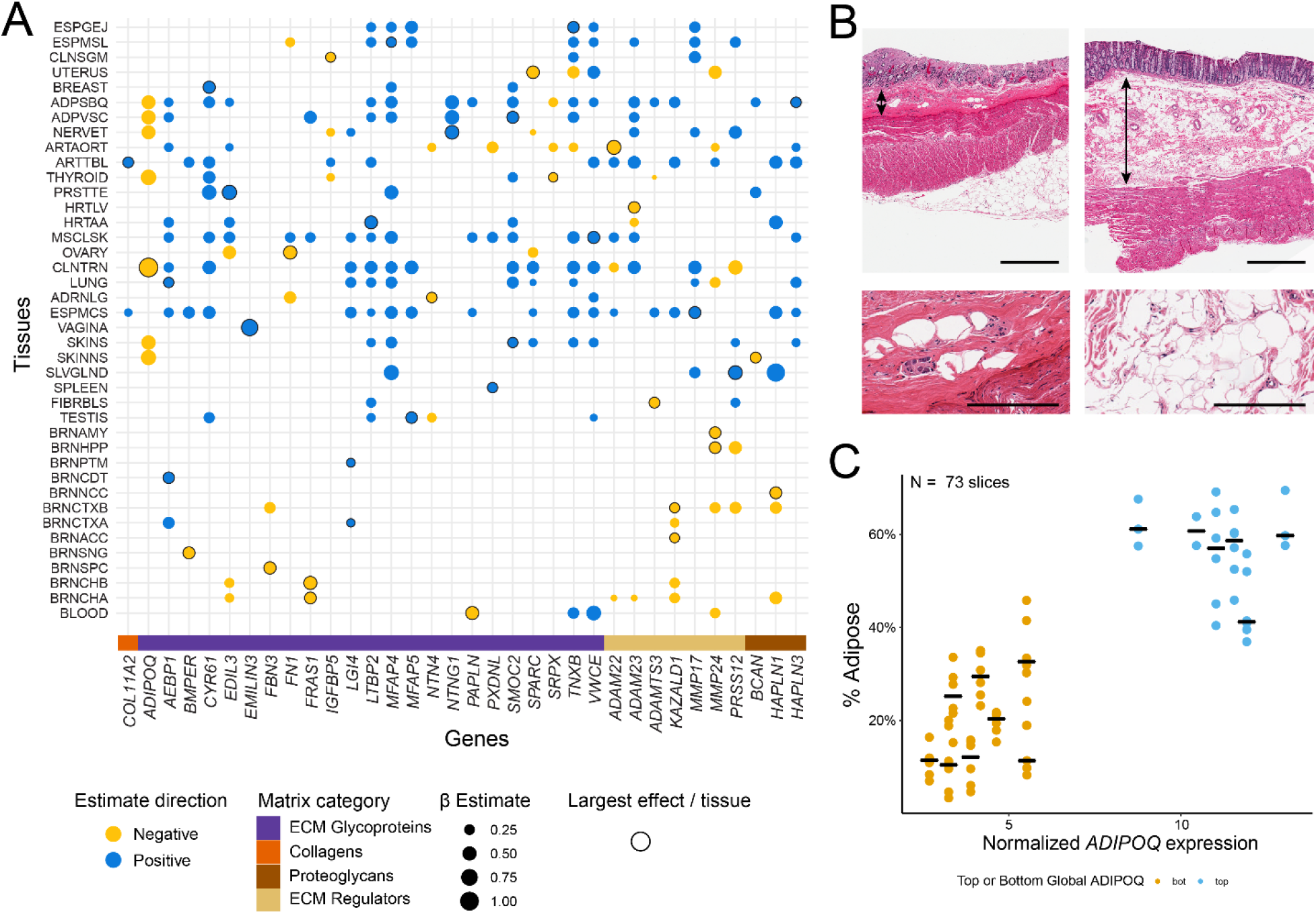
Age-associated transcriptomic changes of the matrisome. A. A plot indicating the 32 genes with the largest change in expression relative to age. The gene with the smallest p-value per tissue is noted along with all other instances of that gene having a β estimate ≥0.15 in other tissues. Tissues with no significant findings were not included. B. Representative images of the transverse colon with varying levels of multi-tissue *ADIPOQ* expression at two different powers. The left sample, GTEX-14C5O, had low levels of *ADIPOQ* and is notable for a thin submucosa (two-headed arrow) with few adipocytes. The right sample, GTEX-1KXAM, had high levels of *ADIPOQ*, with a notable submucosal area containing large numbers of adipocytes. The bars are 1 mm and 200 μm for the low and high power views respectively. C. A scatterplot representing the percent of adipose tissue in the submucosa of the transverse colon for the top 6 and bottom 8 individuals ordered on body wide *ADIPOQ* expression. The datapoints (n = 2 to 6, median of 6, per individual) of multiple measured areas per sample are colored to represent the top (blue) or bottom (yellow) multi-tissue *ADIPOQ* expressors. Bars indicate the median level of submucosal adipocytes for each individual.

One pervasive change across multiple tissues was the reduction in *ADIPOQ* expression levels (7 tissues, mean β = −0.61 sd = 0.23). It was reduced in tissue groups such as adipose tissue, skin tissue, thyroid and tibial nerve, with the largest effect in the transverse colon (linear model; β = −1.109, p = 3.75e-13). It is common for visceral fat to increase with age [17], thus it was unexpected for *ADIPOQ*, a gene associated with adipocytes, to decrease with age in these tissues. To investigate this, we subsetted out individuals ranked on multi-tissue *ADIPOQ* expression into our top (age range: 24-67 years; median: 36.5 years) and bottom (age range: 49-70 years; median: 64 years) *ADIPOQ* groups and evaluated the GTEx transverse colon digitized histologic images.

We observed that, overall, individuals with higher *ADIPOQ* levels had a thicker submucosa with more adipocytes when compared to samples from individuals with lower *ADIPOQ* levels (Fig. 2B). We quantified these differences using ImageJ to determine the percentage of inter-tissue white space, as a surrogate for adiposity, of the submucosal regions. There was a significant correlation between white space and transverse colon normalized *ADIPOQ* levels in the submucosa based on the images taken from the samples with the highest and lowest global ADIPOQ expression levels while controlling for BMI, sex, and age, with subject ID as a random effect (linear mixed model; β = 0.028 p = 5.59e-3; Fig. 2C; Table S2). Age was also significant in this model (β = −0.002, p = 6.25e-3) indicating that the age correlation with reduced submucosal adiposity is not the only factor that contributes to lower submucosal adipose levels.

### Sex differences in the matrisome are driven by sexual dimorphism and Y-chromosome genes

Sexual dimorphism in the transcriptome is characterized by varying cell type composition and sex chromosome expression [18]. Here we attempt to recapitulate these findings and explore them with a focus on matrisome changes. Tissue specific linear models, generated on matrisome gene expression with sex and age as variables identified 325 genes differentially expressed (DE) by sex (222 unique) across 19 tissues (Table S3). As expected, the tissue with the greatest amount of DE genes was breast tissue (n = 134, Figure S4), which was likely driven by the different cell type proportions found in males and females, specifically more adipocytes for males and more epithelial cells for females. *MMP3* was the most commonly DE gene, with significant expression differences in 7 tissues (Fig. 3A).

**Fig 3.**
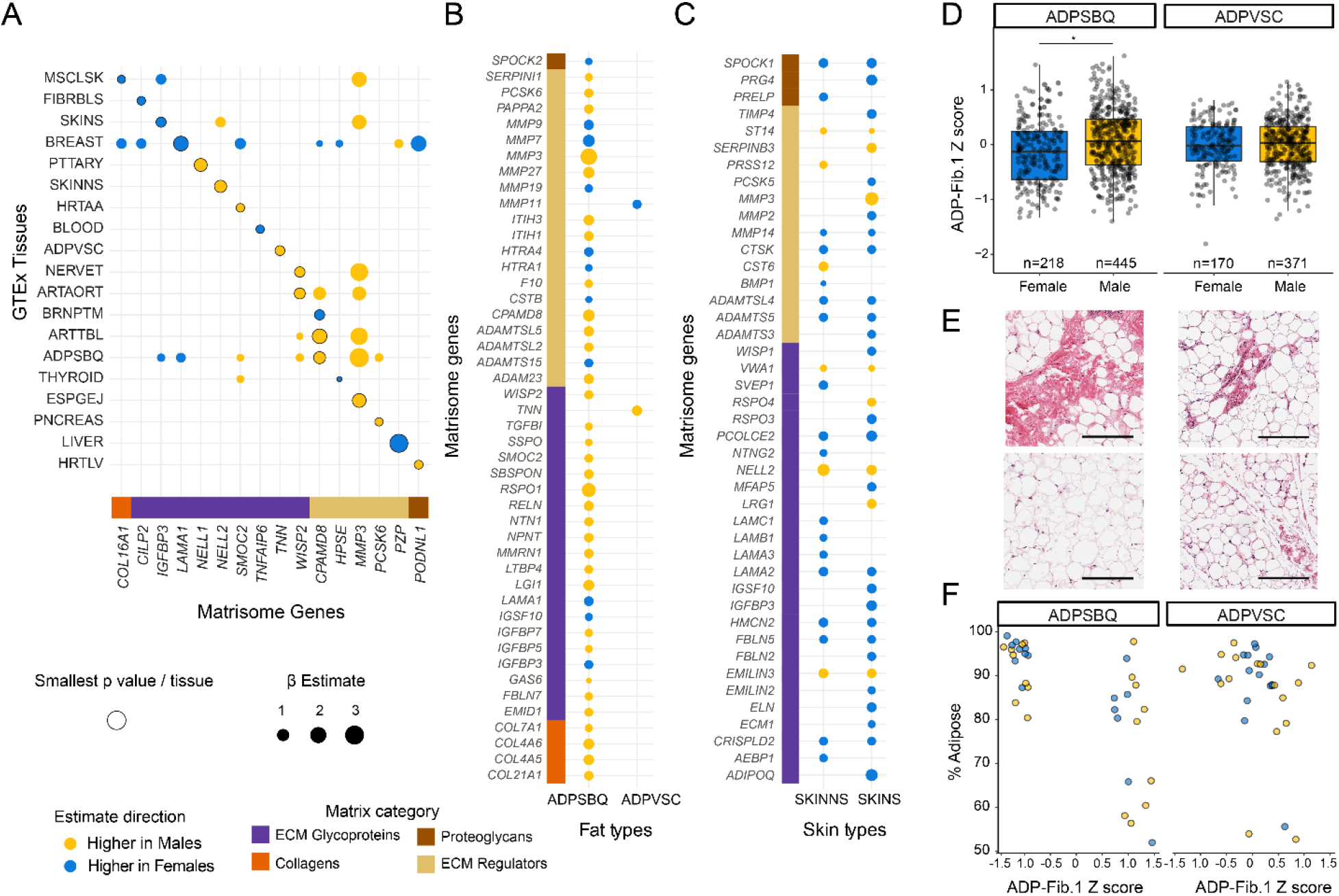
Sex based differences in the matrisome transcriptome. A. A plot of genes with sex DE. The gene with the smallest p-value per tissue is noted along with all instances of that gene having a significant β estimate in other tissues. Tissues with no significant findings were not included. B. The sex-related genes that are DE in ADPSBQ and ADPVSC. Notably, ADPSBQ has 44 DE genes while ADPVSC has only 2. C. The sex-related genes that are DE in SKINNS and SKINS. The size of the dot indicates the β estimate. The tissues are mostly congruent, having 17 shared genes with the same expression directionality. D. A barplot comparing the Z score for the ADPSBQ ADP-Fib.1 gene cluster between males and females for both ADPSBQ and ADPVSC tissues. In ADPSBQ, which has more DE sex genes, cluster ADP-Fib.1 associates with the male sex. This cluster shares 6 genes with the DE gene list in Fig. 3B In ADPVSC, there is no sex difference. E. Adipose samples from two individuals showing differing levels of fibrosis. The top samples are from GTEX-IGN73 and the bottom samples are from GTEX-1K2DA. The left samples are ADPSBQ and the right samples are ADPVSC. The bar is 200 um. F. The median percent adipose tissue from ADPSBQ (n = 34; 16F/18M) and ADPVSC n = 34; 16F/18M) samples, with the ADP-Fib.1 Z-score on the X axis. This data shows that the genes in the ADP-Fib.1 cluster positively correlate with adiposity and negatively correlate with fibrosis in ADPSBQ tissue.

Subcutaneous adipose tissue (ADPSBQ) had the second most sexually DE genes at 44. This contrasted with visceral adipose tissue (ADPVSC) only having 2 DE genes with no genes overlapping between the two adipose tissues (Fig. 3B). Adipose tissue is known to be sexually dimorphic in metabolism and fat deposition [19]. Skin is also characterized by sexual dimorphism [20], a major difference being collagen distribution [21]. Not sun exposed skin (SKINNS) had 24 DE sex genes, while sun exposed skin (SKINS) had 33 genes (Fig. 3C). Unlike the adipose tissue samples, both skin locations shared 14 intersecting genes and directionality, consistent with their having similar functions. ADPSBQ and ADPVSC have more distinct roles in the body [22] and we sought to better understand what drove differential gene expression based on sexual dimorphism.

We clustered co-correlating matrisome genes within ADPSBQ, generating 3 gene clusters. One cluster separated into two negatively correlating sub-clusters termed ADPSBQ fibrosis 1 and 2 (ADP-Fib.1, n = 33 genes; ADP-Fib.2 n = 2 genes) (Table S4). Six genes overlapped between ADP-Fib.1 and the ADPSBQ linear model results, implying this cluster was sexually DE. The expression of ADP-Fib.1 was generally higher in male ADPSBQ samples (β = 0.216, p = 8.96E-06) as determined by a linear model, controlled for BMI on the normalized gene expression from the ADP-Fib.1 cluster (mean = 0, sd = 0.587). This same relationship was not seen in ADPVSC. (Fig. 3D, Table S5).

To better understand this incongruency, we examined the GTEx ADPSBQ histology samples most associated with the extreme ends of the ADP-Fib.1 cluster normalized score, quantifying the amount of adiposity and the inverse amount of fibrosis. This was performed in 73 ADPSBQ and 66 ADPVSC tissue samples from both males (n = 36) and females (n = 32) at the extreme ends of the subcutaneous ADP-Fib.1 Z-scores (Fig. 3E). There was a significant association between the ADP-Fib.1 cluster Z-score and adipose percentage (linear model; 6.87e-5) in a regression model correcting for sex and BMI in ADPSBQ tissue (Fig. 3F). Again, there was no association with ADPVSC (linear model; p = 0.22) using the same model.

### Carcinomas from different tissues have similar matrisome genes changes

We then explored the usefulness of GTEx-based matrisome as a control dataset for cancer stroma studies. To do this, we acquired and compared GTEx and TCGA RNA-seq data for lung, breast, colon, thyroid, and prostate normal tissues and cancers (LUAD, BRCA, COAD, THCA, PRAD) [23]. Using normalized expression values of each matrisome gene, we evaluated the relative rank change of expression between paired normal (TCGA or GTEx) and cancer samples. The relative rank order approach mitigated the expected batch effects problem of working across these datasets. This approach allowed us to identify genes that increased in cancer such as *MMP11* (increase rank by 223 positions) and genes with lower expression in cancer such as *ADAMTS8* (decrease rank by 196 positions) (Fig. 4A).

**Fig 4.**
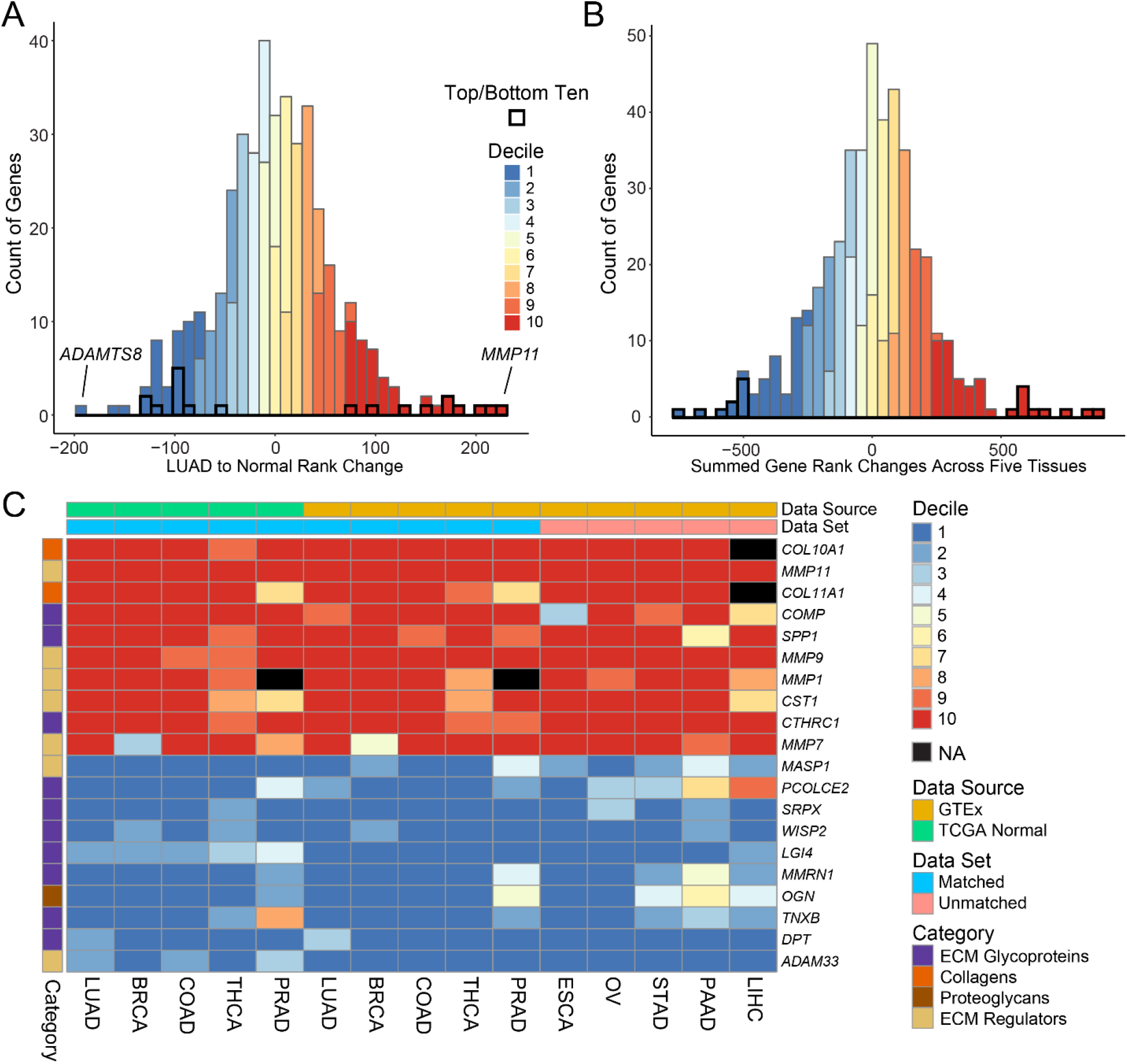
Commonly upregulated and downregulated matrisome genes. A. A histogram of gene rank changes between combined TCGA normal samples (n = 110) and GTEx samples (n = 374) compared to TCGA cancer samples (n = 542) in lung. The histogram is colored on the decile of rank change. B. A histogram of the sum of normal to cancer rank changes for genes across lung, breast, colon thyroid and prostate (n = 443). The histogram is colored on the decile of rank change. C. A heatmap of the top and bottom rank-changed matrisome genes between normal tissue and cancer. The rank change order is split into deciles, (1-1st decile to 10 – 10th decile). NAs are represented by black boxes and indicate no expression in the normal or cancer samples

By summing these normal to cancer rank changes across tissues we generated a list of the top 10 matrisome genes most consistently in the top or bottom decile of rank changes, revealing tissue agnostic cancer expression changes in the extreme ends of the distribution (Fig. 4B, Table S6). These genes were detected equally well between tumor samples matched to TCGA normal (38 of 50 in the 10^th^ decile; 33 of 50 in the 1^st^ decile) or GTEx paired organ samples (39 of 50 in the 10^th^ decile; 41 of 50 in the 1^st^ decile) and represent a fairly consistent set of cancer-associated matrisome changes. We then extended this experiment to esophagus, ovary, stomach, pancreas, and liver carcinomas (ESCA, OV, STAD, PAAD, LIHC) using only paired GTEx normal tissues. In this group, the overexpressed matrisome genes remained consistently in the upper decile (40 of 50), while the lower expressed genes trended in the same direction but were not as consistently in the bottom decile (29 of 50, Fig. 4C).

We confirmed these common matrisome alterations, in six unrelated microarray expression datasets of paired lung, breast, esophagus, and colon carcinomas and normal tissues (Table S7). Differential expression analysis on these datasets demonstrated an average of 8.5 of 10 10^th^ decile genes and 7.8 of 10 1^st^ decile genes were significantly DE in the correct direction across these samples. These findings indicate that GTEx tissue matrisome gene signatures can be a robust control tissue for tumor studies and that a fairly consistent set of up or downregulated matrisome genes are altered among numerous malignancies.

### The matrisome can distinguish idiopathic pulmonary fibrosis from normal and acutely injured lung

Idiopathic pulmonary fibrosis (IPF) is a chronic disease with a progressive decrease in lung function due to increased fibrosis and ECM remodeling [24, 25]. We were interested in determining if gene matrisome data could correlate with the known histopathologic change of IPF and determine if GTEx matrisome data could be used as a control tissue for disease studies. For this, we identified a published gene expression dataset of IPF [26] and integrated this data with the lung GTEx expression data.

We performed clustering analysis of an IPF dataset (Sivakumar *et al.*) containing 46 IPF samples, 8 acute lung injury (ALI) samples and 26 control lung samples along with 20 GTEx samples that we previously indicated were normal or had ventilator-associated injury [12], focusing on the core matrisome genes and ECM regulators [26]. We identified one gene cluster that split into two negatively correlating sub clusters, LUNG-Fib.1 and LUNG-Fib.2 (Figure S5, Table S8). A normalized expression Z-score of genes in each cluster was used to test associations of clusters LUNG-Fib.1 and LUNG-Fib.2 to disease. A linear model, controlling for batch, age, sex, and tobacco use indicated that both ALI and IPF were significantly associated with the cluster LUNG-Fib.1 expression Z-score, with IPF having a larger effect size (linear model; p = 0.017, 3.71e-19; β = 0.47, 1.63; for ALI and IPF respectively). Cluster LUNG-Fib.2 comparatively only correlated with IPF (linear model; p = .0001; β = −0.778).

Hierarchical clustering on the lung samples limited to the genes from clusters LUNG-Fib.1 and LUNG-Fib.2, identified three sample groupings (Fig. 5A). The first grouping contained 35 samples of normal lung, ALI, and ventilator injury from both the GTEx and Sivakumar datasets. It was marked by low expression of gene cluster LUNG-Fib.1 and higher expression of gene cluster LUNG-Fib.2. The second group of 24 samples was a mix of normal, ventilator injury, ALI, and IPF samples, with its expression profile appearing to be an intermediate between groups 1 and 3 for the cluster LUNG-Fib.1 and cluster LUNG-Fib.2. The third grouping contained 40 IPF samples and one normal sample. Clear differences of the LUNG-Fib.1 and LUNG-Fib.2 gene cluster Z-scores can be seen for IPF samples relative to the other injury or non-injury patterns of lung (Fig. 5B). Taken together, this data indicates both that GTEx matrisome data can be usefully integrated with other gene expression datasets and that the matrisome itself can distinguish disease states.

**Fig. 5.**
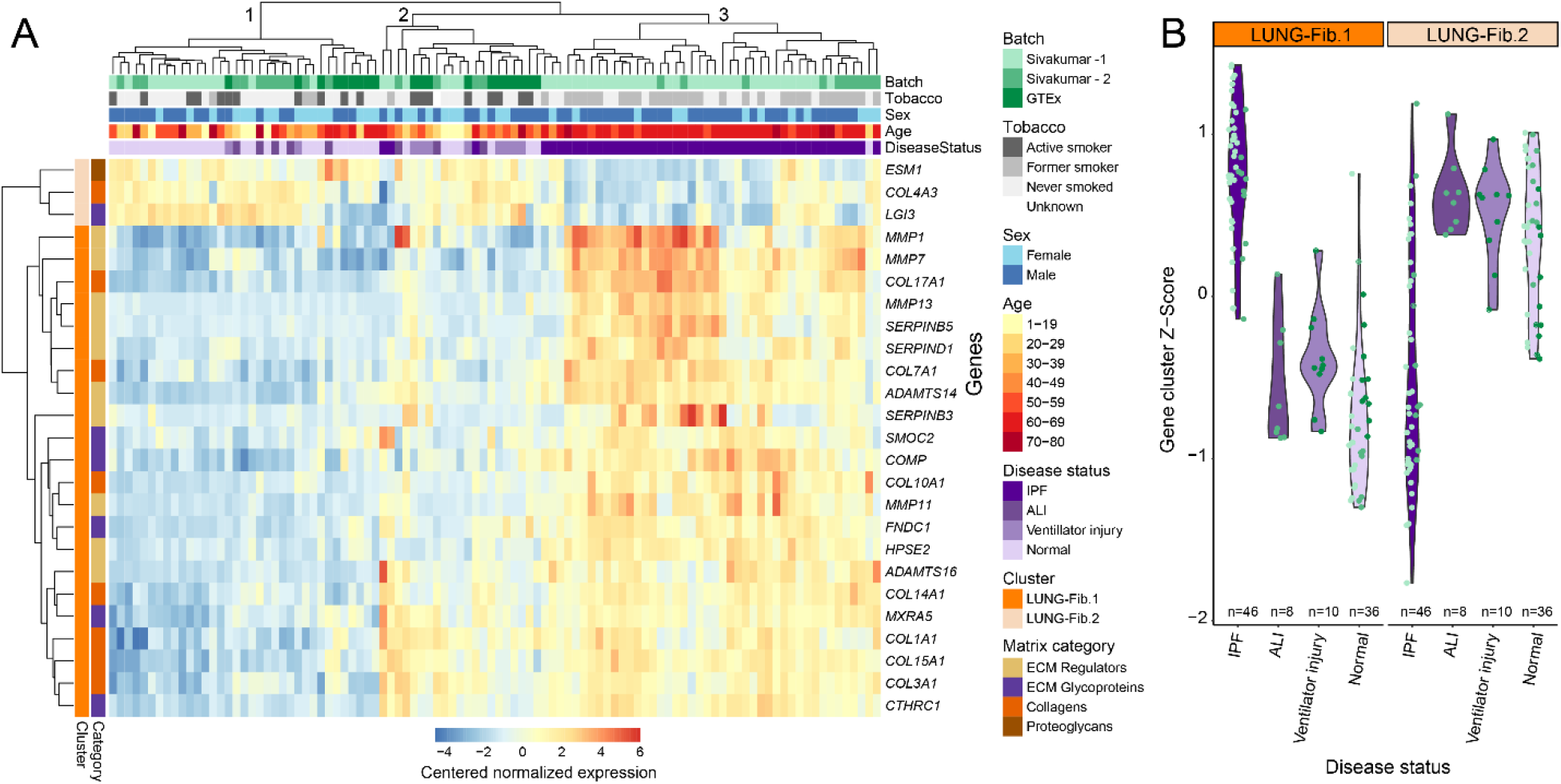
Idiopathic Lung Fibrosis characterization using GTEx. A. A heat map representing the centered normalized expression of matrisome genes in clusters LUNG-Fib.1 or LUNG-Fib.2 between lung samples. The x-axis of the lung samples indicates disease, batch, tobacco usage, sex, and age status of each. Age and being in batch Sivakumar – 2 significantly correlate with IPF samples (logistic regression; p = 0.012; p = 0.029 respectively) B. A violin-sina plot of the normalized score of gene cluster LUNG-Fib.1 and LUNG-Fib.2 comparing IPF, acute lung injury, ventilator injury, and normal lung samples from Sivakumar *et al.* The color of each data point indicates the batch.

## Discussion

Using the GTEx dataset, we have generated a body-wide understanding of the transcriptomic matrisome in health and limited disease states. The large dataset of GTEx, and others like it, may improve our collective understanding of the matrisome by providing complementary data to proteomic and functional studies. As tissue transcriptomic studies are more easily performed and generate deeper datasets than proteomic-based methods, establishing correlation between the two types of data was critical.

We initially validated gene expression as a marker of proteomic levels through the Wang *et al.* LFQ proteomic/gene expression study, and found good correlation with the exception of glycoproteins [8]. However, we believe this poor correlation more likely reflects the challenge of capturing this class by proteomics rather than a weakness of the transcriptomics [10]. Overall, matrisome transcriptomic data clustered similar tissues based on their cell type composition, such as epithelial tissues or smooth muscle tissues and that this matrix gene expression was a reasonable surrogate for proteomic expression (Fig. 1C).

The second proteomic study, 10-plex TMT, was performed on GTEx tissues and was expected to correlate well with the transcriptomic data [8]. It did not perform as well. One of the challenges of a 10-plex TMT when working with diverse samples, is having a control sample on each TMT that contains every peptide needed to normalize proteins for all tissues. The TMT approach pooled reference samples across 32 organs, which diluted out rare/organ-specific proteins (~1/32^rd^ of the peptide mix) reducing capture of these peptides and decreasing the organ specificity of control peptides.

While other studies have investigated aging throughout all GTEx gene expression on earlier versions of GTEx data, we focused on just matrisome gene aging correlations [27]. Melé et al. generated a list of 47 DE matrisome genes due to aging across all GTEx tissues. With a larger dataset, we identified 440 significantly changed matrisome genes analyzed individually across multiple tissues, capturing all but two, *MMP12* and *LEPRE1*, of the genes in their dataset. However, of the 45 shared genes, only 60% shared global directionality. These discrepancies are most likely due to their analysis being on all genes, resulting in a stricter p-value threshold, and their usage of tissues as a random effect in their model.

Our findings reveal that there are tissue specific age related changes in the matrisome. Some have bodywide directionality such as a decrease in *ADIPOQ* and an increase in *MFAP4*, a possible biomarker associated with cardiovascular conditions and liver fibrosis [28, 29]. Other genes did not always share body-wide directionality such as *TNXB* and *PRSS12* which implies genomic tissue specific matrisome changes with age. Using GTEx’s histology data we were able to elucidate an association between *ADIPOQ*, a gene decreasing with age in multiple tissues, and loss of fat in the CLNTRN sub-mucosal space. Our findings are inconsistent with other literature stating that inter-mucosal fat increases with BMI [30] and does not decrease with age [31]. However our data differs from theirs in several regards. Wada *et al.* studied the fat using liquid droplets, and Mesa *et al.* looked at submucosal thickness as a measure of fat in comparison to our approach which evaluated the percentage of submucosal space by histology.

Our analysis has also generated a list of matrisome specific differences between the sexes. Breast tissue, due to its sexual dimorphism, was logically the tissue with the most differentially expressed genes. The most prevalent sex differentially expressed gene was *MMP3*. Previous work has shown that the *MMP3* protein is elevated in male serum or plasma when compared to females [32, 33]. However, our finding appears to be the first description of *MMP3* differences across multiple tissues and at the gene level. The fat and skin data had striking differences between tissues and sex (Fig 3C and 3D). ADPSBQ had 75% of 44 DE genes having higher expression in males, while ADPVSC had only 2 DE genes. Conversely, SKINS and SKINNS had ~75% of DE genes (33 and 24, respectively) having higher expression in females. There are important functional differences between subcutaneous and visceral adipose tissue characterized by sex-specific and hormone-based differences [22, 34], while the skin sampling locations (suprapubic and lower leg) were expected to be more similar. Sex DE matrisome genes were investigated previously using GTEx v6 [18]. That study found 242 unique sex DE matrisome genes, on a per tissue basis, of which 150 overlapped with our findings. Differences likely result from methodological approaches, GTEx versions, and multiple testing corrections.

Integrating GTEx and TCGA allowed us to explore the cancer matrisome. Focusing on carcinomas, a combination of GTEx and TCGA decribed a list of genes consistently upregulated or downregulated, even across tumor types for which the TCGA has a paucity of adjacent normal tissues. The top gene, *COL10A1* is a well-known marker of solid tumors [35] and *COL11A1*, the third highest gene, is a marker of cancer-associated fibroblasts [36]. Among downregulated genes, the *ADAM33* promoter is silenced in aggressive breast cancer [37, 38]. *DPT* downregulation in oral cancer is associated with increased metastasis [39]. These rank changes, validated in unrelated gene microarray datsets, show the robustness of this gene set.

Using the high variance matrisome gene clusters, we were also able to parse out associated DE genes that may play roles in the pathology of IPF. A prior manuscript evaluated global gene changes in IPF describing *COL1A1* and *MMP7* changes in IPF, similar to our findings, along with several genes (*LAMB3, LAMC2, FN1*, and *TNC*) not uncovered in our project [25]. Importantly, the GTEx samples overlapped appropriately with the non-IPF samples of the Sivakumar study, indicating their value as a control tissue source for other matrisome studies across a range of disease states.

This study has several limitations. First, GTEx data is skewed, with a bias for male subjects (n = 653 males, n = 327 females) and a bias for older individuals (mean = 52.76 years, median = 55 years, standard deviation = 12.91 years). The lack of younger individuals likely impacts on the aging analysis of matrix genes that reach steady-state later in life [40]. Contamination from highly expressed genes in GTEx, is known, and could impact on specific expression patterns [11]. There were also limitations due to GTEx's available histology. Not all individuals had histology samples of the tissues we interrogated, and for some, the tissue orientation precluded image analysis. While it would have been interesting to evaluate the tumor matrisome in metastases, TCGA had many fewer samples of this type.

In conclusion, patterns of matrisome gene expression approximate the measured values of matrix proteins. As a result, the GTEx database is a robust resource for transcriptomic matrisome gene expression to explore matrix genes variability across tissues, age and sex. Further, in two examples, it demonstrated usefulness as a control tissue source for disease studies and can be used in this capacity to augment further investigations into the human matrisome.

## Experimental Procedures

### Retrieval and correlation of Label Free Quantification / Genomic datasets

We acquired protein and protein-coding gene data from Wang *et al.* [8] found in their supplementary table 1 and supplementary table 2 respectively. The proteins were analyzed by mass spectrometry using identification and intensity based absolute quantification (iBAQ) [41]. Methods of RNA extraction of the tissues, including library preparation and sequencing are described in Uhlén *et al.* [42], our analysis used their fragments per kilobase of transcript per million mapped reads (FPKM) data.

All analyses were completed in R v.4.0.2 unless stated otherwise. For each tissue, we linked their protein iBAQ and gene FPKM data together. The data was filtered down to matrisome core and regulatory protein-gene pairs that had non-0 values in their gene and protein columns. Using base R, Spearman’s rank correlations were generated for the following comparisons: correlations between genes and proteins within each tissue; and a distribution of correlations for each tissue, randomly sampling N non-0 protein-gene pairs (without replacement) 10,000 times using dplyr 1.0.2. The N of samples is the number of viable matrisome genes for each tissue. This process was repeated for each matrisome category.

### Retrieval and correlation of GTEx proteomic-transcriptomic data

Protein data and gene data were retrieved from Supplementary Table 2D and Supplementary Table 3A from Jiang *et al.* [9]. The protein data used was “Cleaned relative protein abundances in log2 scale” and the gene data was “Cleaned and normalized RNA log2 (TPM) expression for all protein-coding genes across matched samples”. We also acquired their gene across tissue correlation data from Supplementary Table 4A.

From the proteomic and transcriptomic data, samples from individuals with proteomic and transcriptomic data were selected for the analysis. The protein and gene samples were joined and filtered, requiring the protein-gene pairs to have non-0 and non-NA values in each column. We then generated a Spearman’s rank correlation for each sample pair on all genes, and the core and regulatory matrisome.

Using the body wide gene tissue correlations from Jiang *et al.* (Supplementary Table 4A; 1-s2.0-S0092867420310783-mmc5.xlsx, sheet 2) each protein-gene pair correlation was labeled with its matrisome category or as “Non-Matrisome”. Each matrisome category’s correlations were compared against the correlation of all other genes using a Mann-Whitney U test. The resulting p-values were corrected for multiple tests using Holm’s method, and the data was presented using ggplot2 (v.3.3.2).

### Retrieval and Processing of GTEx Bulk Sequencing and Phenotype Data

The gene read counts of the RNA-Seq GTEx version 8 data set (GTEx_Analysis_2017-06-05_v8_RNASeQCv1.1.9_gene_reads.gct.gz) were downloaded from the GTEx Portal (https://gtexportal.org/home/datasets), along with the de-identified sample and subject annotations (GTEx_Analysis_v8_Annotations_SampleAttributesDS.txt, GTEx_Analysis_v8_Annotations_SubjectPhenotypesDS.txt). GTEx v8 median TPM for each tissue was downloaded through the GTEx portal (GTEx_Analysis_2017-06-05_v8_RNASeQCv1.1.9_gene_median_tpm.gct.gz). Numeric ages of the GTEx subjects were acquired through dbGap with approval.

For the median tissue expression, raw read counts were normalized together using the VST feature in DESeq2 (v.1.22.1) in R version (v.3.6.117). This method incorporates estimated size factors based on the median-ratio method, and transformed by the dispersion-mean relationship.

The median VST normalized read counts for core matrisome and ECM regulator genes were used to develop a tissue median expression profile. The median expression of all separate brain tissues were used to form the median BRAIN tissue. A clustered heatmap of tissues were generated using a Kendall’s τ correlation matrix. The heatmap was plotted using Pheatmap (v.1.0.12) and the tissues were clustered using a 1 – Kendall’s τ correlation matrix based on their Euclidian distance. Matrisome genes were labelled with their respective categories, and each category was separately summed for each tissue, and then divided by the total TPMs within each tissue. These values were plotted using ggplot2.

For the age and sex analysis raw read counts, by tissue type, were filtered using the filterByExpr function from edgeR (v.3.32.1) [19910308] before being normalized by the voom feature [24485249] in limma (v.3.46.0) [25605792] in R version (v.4.0.2). This method generates log2-counts per million, and is suited to eliminate the mean-variance relationship found in RNA-seq data using observational-level weights. GTEx tissues were filtered to only core matrisome and ECM regulator genes. Independently, each of these genes were modeled as a linear function of age and sex using limma’s lmFit function. The results for all tissues were collated together and had a Holm’s correction applied to their p-values. Only estimates with a >0.05 Holm’s corrected p-value were used for further analysis. The data was plotted using ggplot2, limited to genes with smallest p-values for the given covariate in each tissue.

### Sex based differences of Adipose Tissue and Transverse Colon

For both ADPSQB and ADPVSC, previously VST normalized genes were required to have a mean normalized read count >5 and an across sample variance >1.5 for inclusion in the cluster analysis. The filtered genes were clustered using hierarchal clustering on a distance generated by 1−Kendall’s rank-correlation coefficient. A τ critical value was calculated based on the number of samples and genes expressed. The correlation-based dendrogram was cut to produce gene clusters with an average cluster correlations of at least the τ critical value.

Normalized expression scores were calculated by subtracting the mean expression and dividing by the median absolute deviation of the expression values for each gene across all samples within a given tissue. The equation is as follows, where *x* is the VST normalized expression of gene *j* in sample *i*, *t*(*i*) is the tissue type for sample *i*, and *J* is the number of genes in a given cluster.

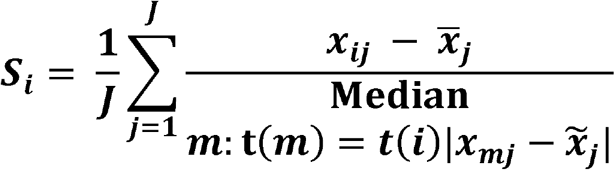

Using the previously generated ADP-Fib.1 cluster score, 5 males and 5 females were selected from the highest and the lowest ADP-Fib.1 scores for both ADPSBQ and ADPVSC. These samples were then de-identified and randomized to prevent biased ImageJ scoring.

Previously VST normalized CLNTRN, CLNSGM, ADPSBQ, and ADPVSC samples were selected on the requirement that all four tissues were available for a given individual. This RNA-seq data was limited to only the *ADIPOQ* gene, and normalized reads were averaged across the tissues generating the multi-tissue *ADIPOQ* score. The individuals were sorted on the multi-tissue *ADIPOQ*, and the top 6 individuals and the bottom 8 individuals were selected to have their transverse colon analyzed using ImageJ.

The percent adipose content of each sample was determined using ImageJ 2.0.0 (FIJI) [43]. For the transverse colon, only samples with submucosal space present were evaluated. The ImageJ freehand selection tool, captured the area of the submucosal space, excluding veins, white space, and muscle. For ADPSBQ and ADPVSC, the fat area was captured, avoiding white space and vasculature. The images were transformed to greyscale, a threshold was applied (range 199 to 255 light) and the percent of captured area being either adipose (white space) or fibrosis (colored space) was calculated.

Two linear models were applied on both ADPSQB and ADPVSC. The first model used BMI and sex as covariates for the ADP-Fib.1 score. The second model used ADP-Fib.1 score, sex, and BMI as covariates for percent adipose of the tissue samples. Individuals with more than one sample per tissue had the average of the percent adipose tissue used as their value.

For the transverse colon model, multi-tissue normalized *ADIPOQ* score, age, and BMI were used as covariates on percent adipose tissue in transverse colon samples. As there were multiple tissues slices per person, subject ID was used as a random effect in the model. The R package lme4 (v.1.1-26) was used to generate the linear mixed model. In the model equation (below), *β*_1_ is the *ADIPOQ* Score for sample *i*, *β*_2_is the age of individual *j*, *β*_3_ is the sex of individual *j*, *β*_4_ is the BMI of individual *j*, *u*_*n*(*i*)_ is the random effect of the individual for sample *i*, and *ϵ*_*i*_ is the error term for said sample.

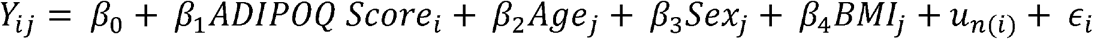

### Generating tissue-agnostic rank changes between normal and cancer samples

GTEx data and TCGA data was acquired through the R package TCGAbiolinks v.2.18.0. The normal GTEx and normal TCGA tissues downloaded were colon (n = 376 GTEx [CLNTRN], n = 51 TCGA), prostate (n = 119 GTEx, n = 52 TCGA), breast (n = 92 GTEx [female], n = 111 TCGA), thyroid (n = 361 GTEx, n = 59 TCGA), lung (n = 374 GTEx, n = 110 TCGA), ovary (n = 108 GTEx), stomach (n = 204 GTEx), pancreas (n = 197 GTEx), esophagus (n = 331 GTEx [ESPMCS]), and liver (n = 136 GTEx), with their respective TCGA cancers being COAD, PRAD, BRCA, THCA, LUAD, ESCA, OV, STAD, PAAD, LIHC. Using TCGA histological types, COAD was limited to colon adenocarcinoma (n = 426), prostate to prostate adenocarcinoma acinar type (n = 489), breast to infiltrating ductal carcinoma (n = 788), thyroid to thyroid papillary carcinoma (classical) (n = 357), ovary to serous cystadenocarcinoma (n = 430), stomach to both stomach adenocarcinoma and stomach intestinal adenocarcinoma (n = 306), pancreas to pancreas-adenocarcinoma ductal type (n = 147), and liver to hepatocellular carcinoma (n = 356). Lung adenocarcinoma (n = 542) and esophageal carcinoma (n = 184) did not require further filtration. Normal TCGA samples for esophagus, ovary, stomach, liver, and pancreas were excluded.

Separately, all tissues and their datasets, GTEx normal (GTEx-N), TCGA normal (TCGA-N), and TCGA cancer (TCGA-C) were filtered down to genes with an across sample sum >5 raw counts (N_min_/N_max_ = 49,463 – 56,114 genes, liver and stomach respectively) and normalized using the DESeq2 VST function. After normalization, genes were further filtered to include only those with a sample mean >5 normalized counts (N_min_/N_max_ = 20,881 - 25,075 genes, liver and thyroid respectively) generating a total gene whitelist for the analysis. This allowed inclusion of genes that were lowly expressed in normal tissue, but had higher expression in cancer tissue.

The above analysis was replicated after normalization steps and whitelist formation, filtering down to core matrisome and ECM regulatory genes (N_min_/N_max_ = 353-410, liver and lung respectively). GTEx-N, TCGA-N, and TCGA-C datasets were combined together and quantile normalized using the package preprocessCore (v.1.52.0) (https://github.com/bmbolstad/preprocessCore), before being separated again. The quantile normalized mean counts for genes in each dataset were calculated. An average was taken between the GTEx-N’s and TCGA-N’s mean quantile normalized counts to form a joined normal mean (Joined-N; Figure S6). Each gene list was ordered on their mean expression, generating for each tissue a list of ranked matrisome genes from GTEx-N, TCGA-N, TCGA-C, and Joined-N used to calculate a numerical rank change for each gene, between normal and cancer tissues (Joined-N compared to TCGA-C, Figure S7). For esophagus, ovary, stomach, liver, and pancreas, GTEx-N was used in place of Joined-N.

Relative rank changes of genes between TCGA-C and Joined-N were determined for lung, breast, colon, thyroid, and prostate tissue. This was replicated between TCGA-C and GTEx-N for esophagus, ovary, stomach, liver, and pancreas. The TCGA-C to Joined-N rank changes per gene were summed across the five tissues to establish a multi-tissue normal-to-cancer rank change. This highlighted genes that consistently increased or decreased across cancer samples, while lowering the effect of tissue specific changes or changes inadvertently caused by the rank change of other genes. The rank changes were segmented into deciles and the top and bottom 10 genes from the multi-tissue rank change were subsetted out and plotted using Pheatmap.

Six separate microarray studies were used to recapitulate the matrisome cancer findings. Two lung datasets (GSE31210, GSE19188), two breast datasets (GSE15852, GSE109169), an esophagus dataset (GSE161533), and a colon dataset (GSE44076) were obtained through GEOquery 2.58.0. DE genes were discovered using the limma 3.46.0 R package’s eBayes function [44] with Holm’s adjustment for p-value correction. For each gene, if ≥50% of probes for a given gene were called as significantly DE, the gene was considered DE.

### Acquisition, processing, and gene variance clustering of IPF data

From GEO, the design, gene annotations, and raw counts for GSE134692 were downloaded, including their IPF, ALI, and normal samples (n = 46, 8, and 26 respectively) [26]. The data was processed as the adipose tissue above, however the variance filter was a >2 variance in this analysis. This returned two sub-clusters of high variance matrisome genes for the IPF dataset.

Histological images of GTEx lung samples were previously categorized as normal or ventilator injured by a lung pathologist [12]. From this group, 10 samples were randomly selected from both the ventilator injury samples and the normal samples.

GTEx lung samples and samples from Sivakumar *et al.* were combined into one DESeqDataSet the design including both disease status (IPF, ALI, Ventilator Injury, and normal) and Batch (Sivakumar-1, Sivakumar-2, and GTEx). Sivakumar *et al.* mentions they correct for these batches, but do not give them further description. The data was normalized using DESeq2’s VST transformation. After transformation, the data was filtered to only contain the high variance IPF gene clusters (n = 25) before being centered on the average gene expression across samples. The samples and genes were clustered using Euclidian distancing and graphed using the Pheatmap package. All analysis method information has been deposited at GitHub https://github.com/tnieuwe/Matrisome_GTEx_analysis.

## Author Contributions

Conceptualization: TON, AZR, MKH; Analysis: TON; MNM; Writing: TON, MKH; Reviewing & editing: all authors.

## Acknowledgements

This work was supported by the National Institutes of Health grants 1R01HL137811 (MKH), R01GM130564 (MKH), R01GM139928 (MNM), and the University of Rochester CTSA award number UL1TR002001 (MNM).

## Abbreviations

GTEx: Genotype Tissue Expression Project
TCGA: The Cancer Genome Atlas
IPF: Idiopathic pulmonary Fibrosis
LFQ: Label free quantification
TMT: Tandem mass tag
iBAQ: Intensity-based absolute quantification
sd: standard deviation
LCL: EBV-transformed Lymphocytes
DE: Differential Expression
ADPSBQ: Subcutaneous adipose tissue
ADPVSC: Visceral adipose tissue
SKINS: Sun exposed skin
SKINSS: Not sun exposed skin
ADP-Fib.1: A subcluster of co-expressed genes found in ADPSBQ
ADP-Fib.2: A subcluster of co-expressed genes found in ADPSBQ
LUAD: TCGA Lung Adenocarcinoma
BRCA: TCGA Breast cancer
COAD: TCGA colon
THCA: TCGA thyroid
PRAD: TCGA prostate
ESCA: TCGA esophagus
OV: TCGA ovary
STAD: TCGA stomach
PAAD: TCGA pancreas
LIHC: TCGA liver
ALI: Acute lung injury
LUNG-Fib.1: A subcluster of high variance co-expressed genes found in Sivakumar et al.
LUNG-Fib.2: A subcluster of high variance co-expressed genes found in Sivakumar et al.

## Supplementary Material

Tables:

Table S1. Genes that significantly correlated with age in a linear model

Table S2: Mixed linear model results of the CLNTRN submucosal fat analysis

Table S3: Genes that significantly correlated with sex in a linear model

Table S4: ADPSBQ high-variance gene clusters

Table S5: Linear model analysis of sex adipose differences

Table S6: Rank sum changes from Joined-N to cancer

Table S7: Recapitulate cancer results in microarray

Table S8: High-variance lung idiopathic lung fibrosis gene clusters

## Supplemental Figures

**Figure S1.**
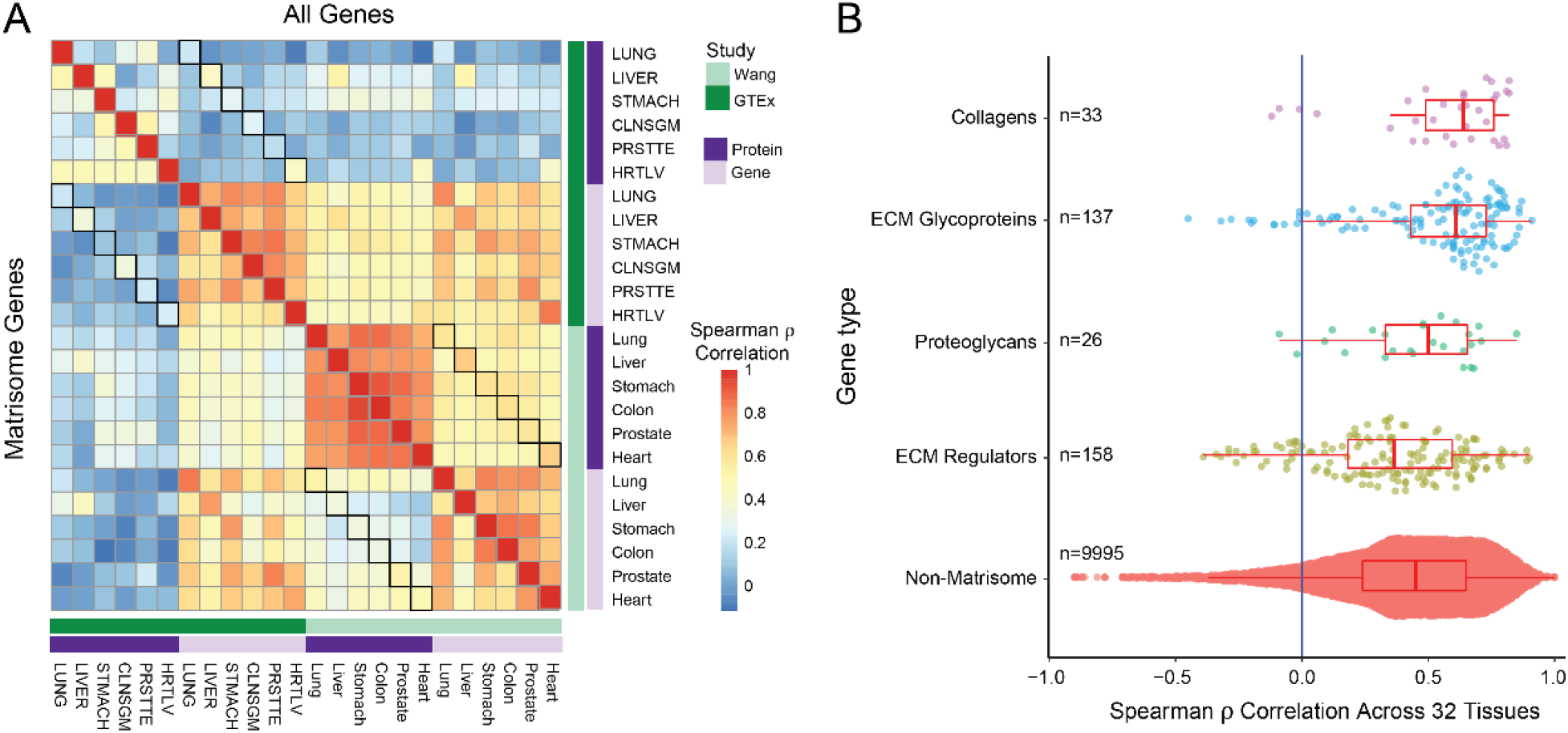
GTEx proteomic data poorly correlates with GTEx transcriptome data while Wang et al. proteomic data correlates well with both GTEx and Wang transcriptomic data. A. A Spearman’s correlation heatmap of six tissues shared across GTEx and Wang *et al.* studies between proteomic and genomic datasets, all genes on the top portion and matrisome genes on the bottom. Using randomly selected representative samples of GTEx tissues, we correlated the shared genes between the two datasets for each tissue (all genes min = −0.193 max = 0.914, matrisome genes min = −0.258, max = 0.942). Black box outlines indicate key correlations. Overall, the GTEx proteomic data correlates the worst with the other datasets. B. A sina and boxplot of different matrisome categories correlations across tissues in the GTEx proteomic and genomic dataset. Collagens and glycoproteins significantly outperformed non-matrisome genes, while ECM Regulators underperformed (Mann-Whitney; p = 2.71e-03, 1.32e-06, and 6.03e-03 respectively). Proteoglycans did not significantly over or under perform non-matrisome genes.

**Figure S2.**
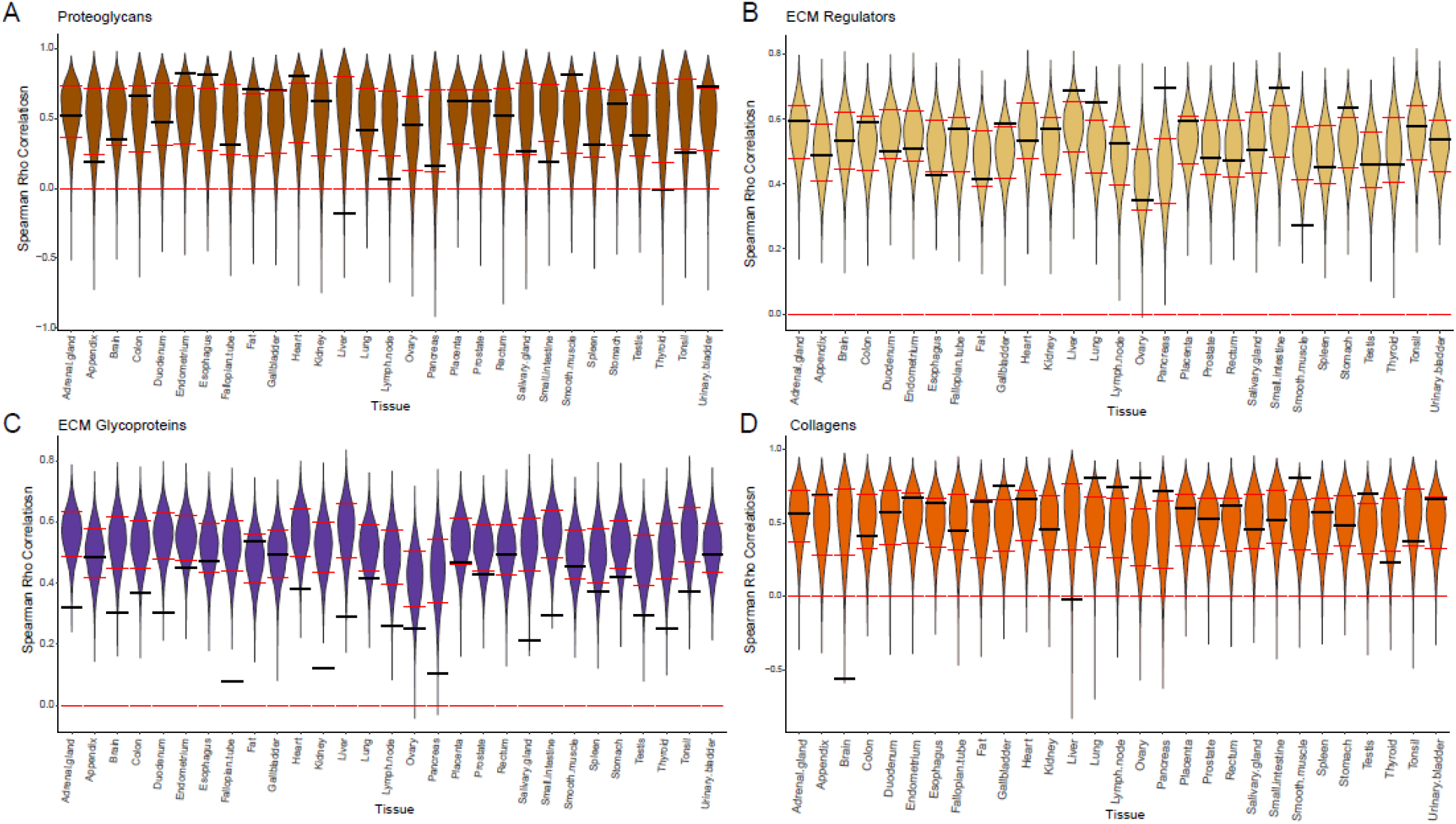
Matrisome category base violin plots of 10,000 permuted Spearman’s correlations and each tissue’s matrisome correlation. Three classes, proteoglycans, ECM regulators and collagens, had strong gene/protein correlations. ECM glycoproteins had overall poor gene/protein correlations.

**Figure S3.**
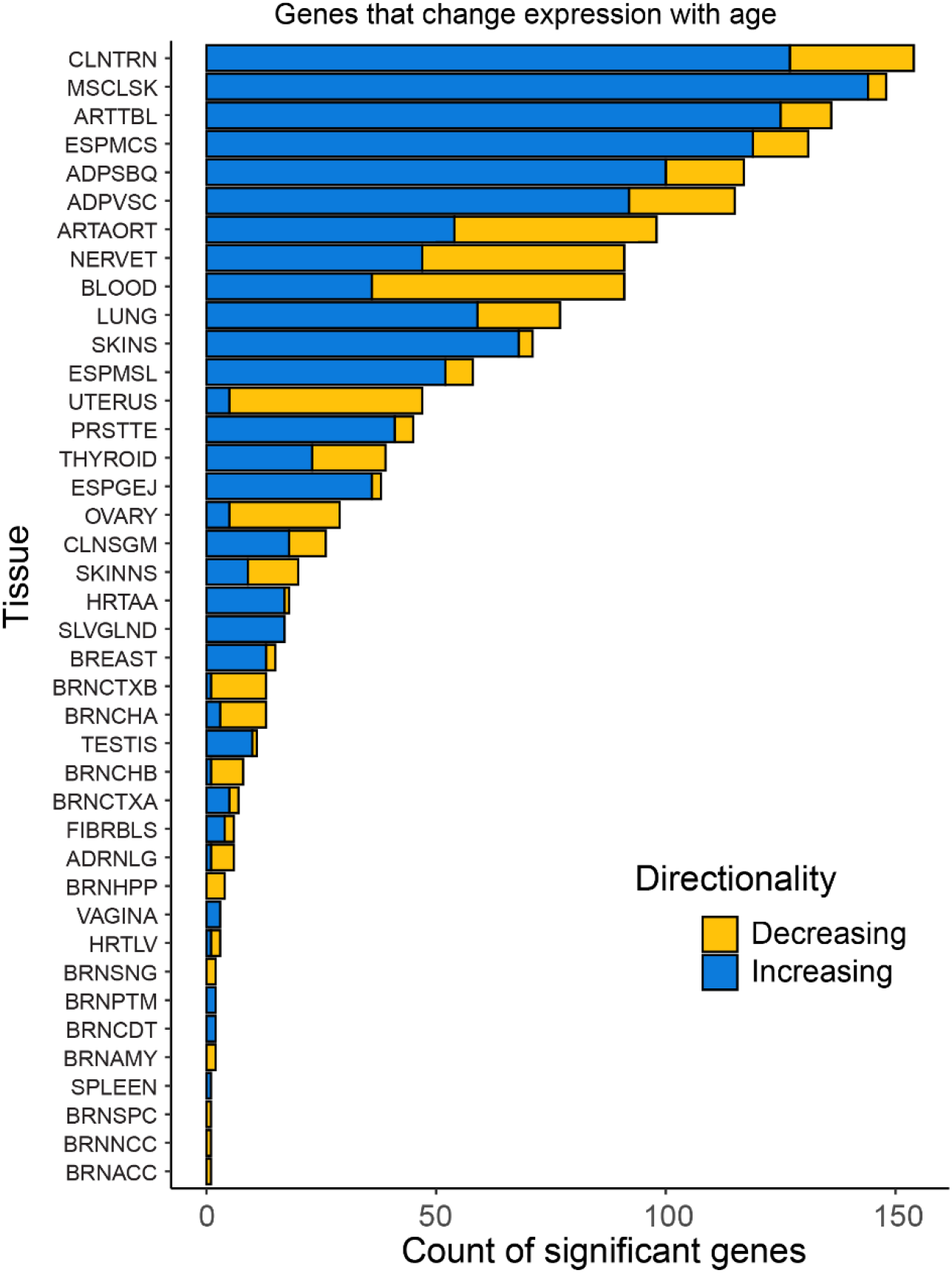
A barplot count of matrisome genes for each GTEx tissue that significantly change with age based on a linear model controlling for sex.

**Figure S4.**
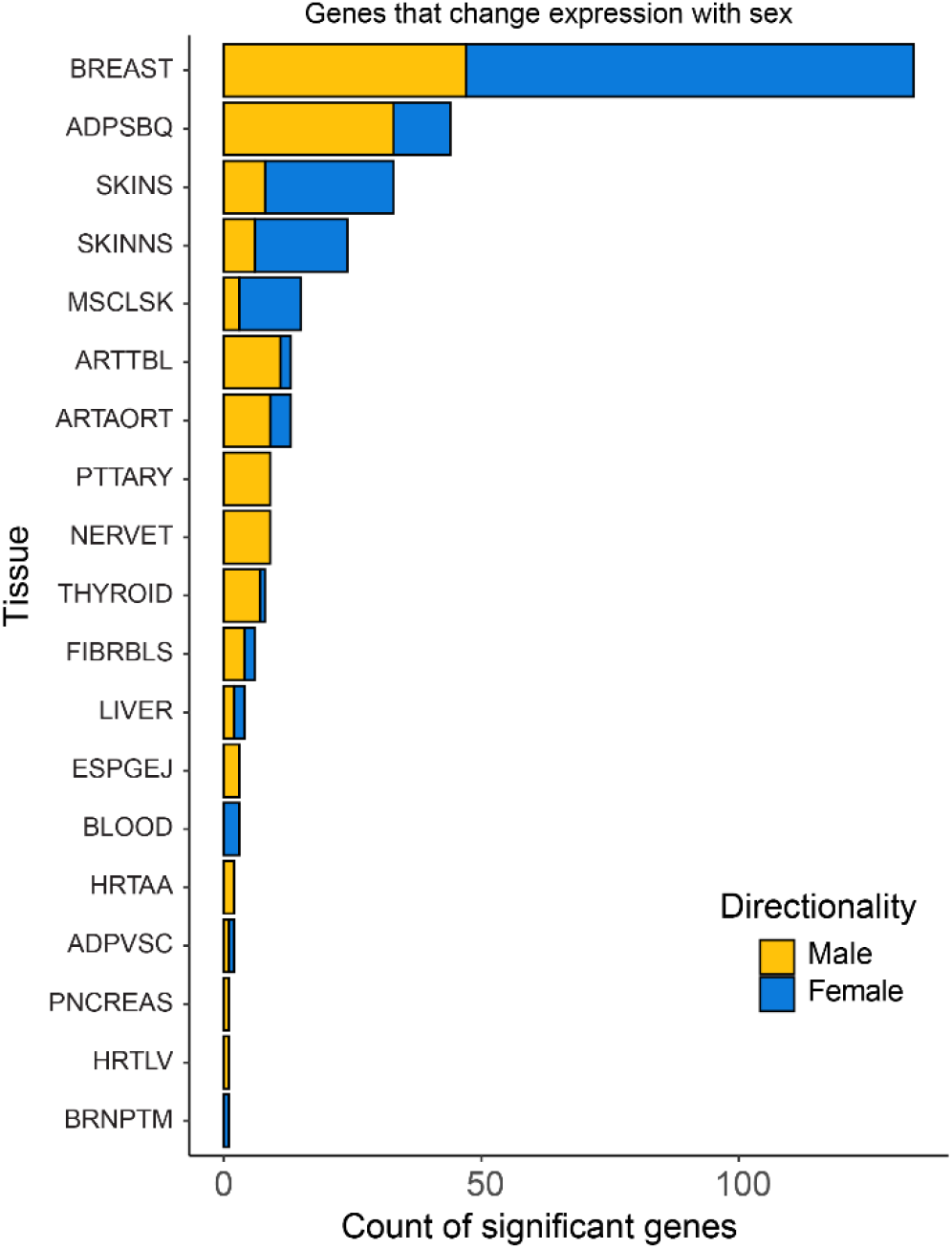
A barplot count of matrisome genes for each GTEx tissue that significantly change with sex based on a linear model controlling for age.

**Figure S5:**
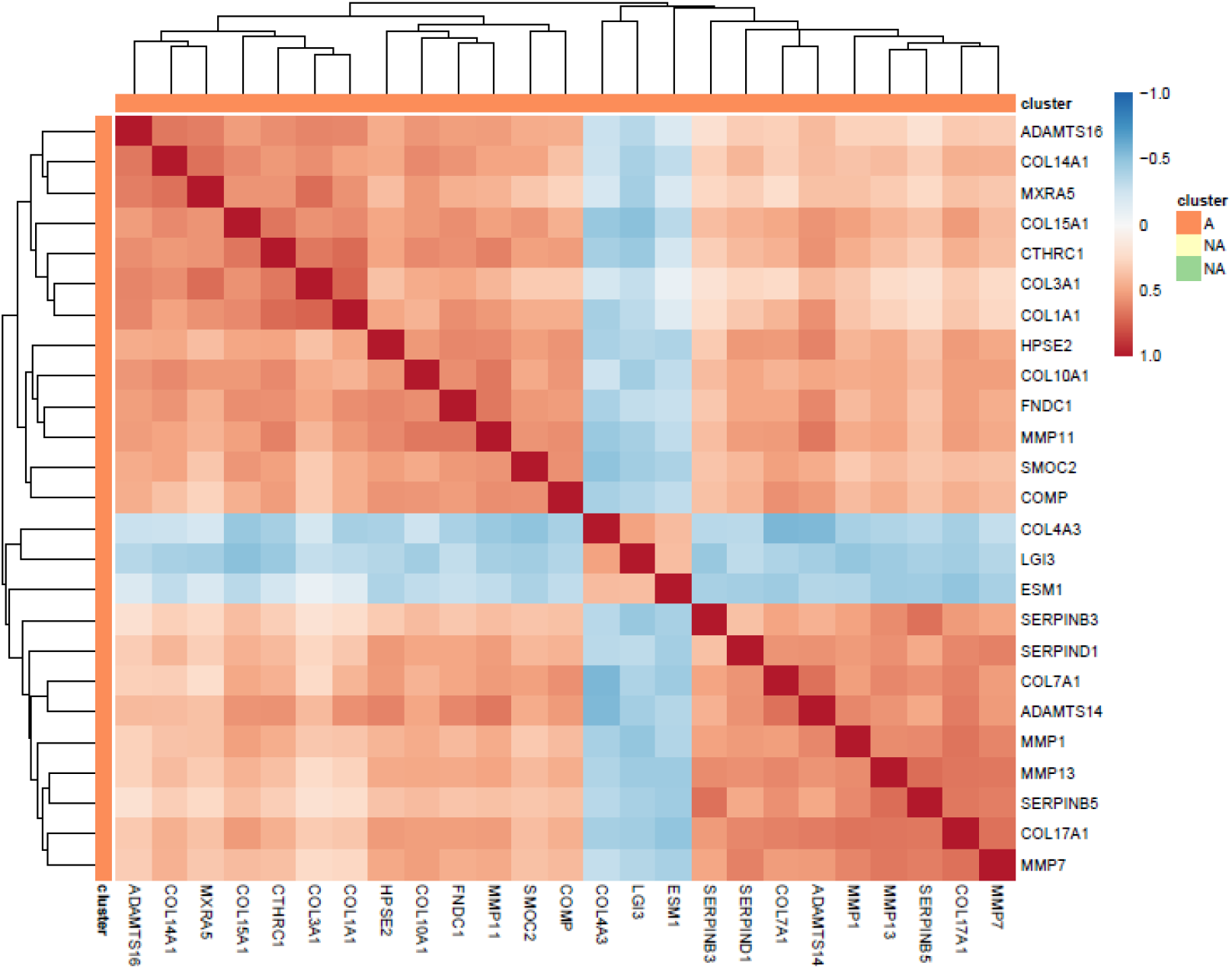
A heat map of the Kendall’s τ correlation matrix between high variance matrisome genes in the GSE134692 data set that includes IPF, ALI, and normal lung tissue.

**Figure S6:**
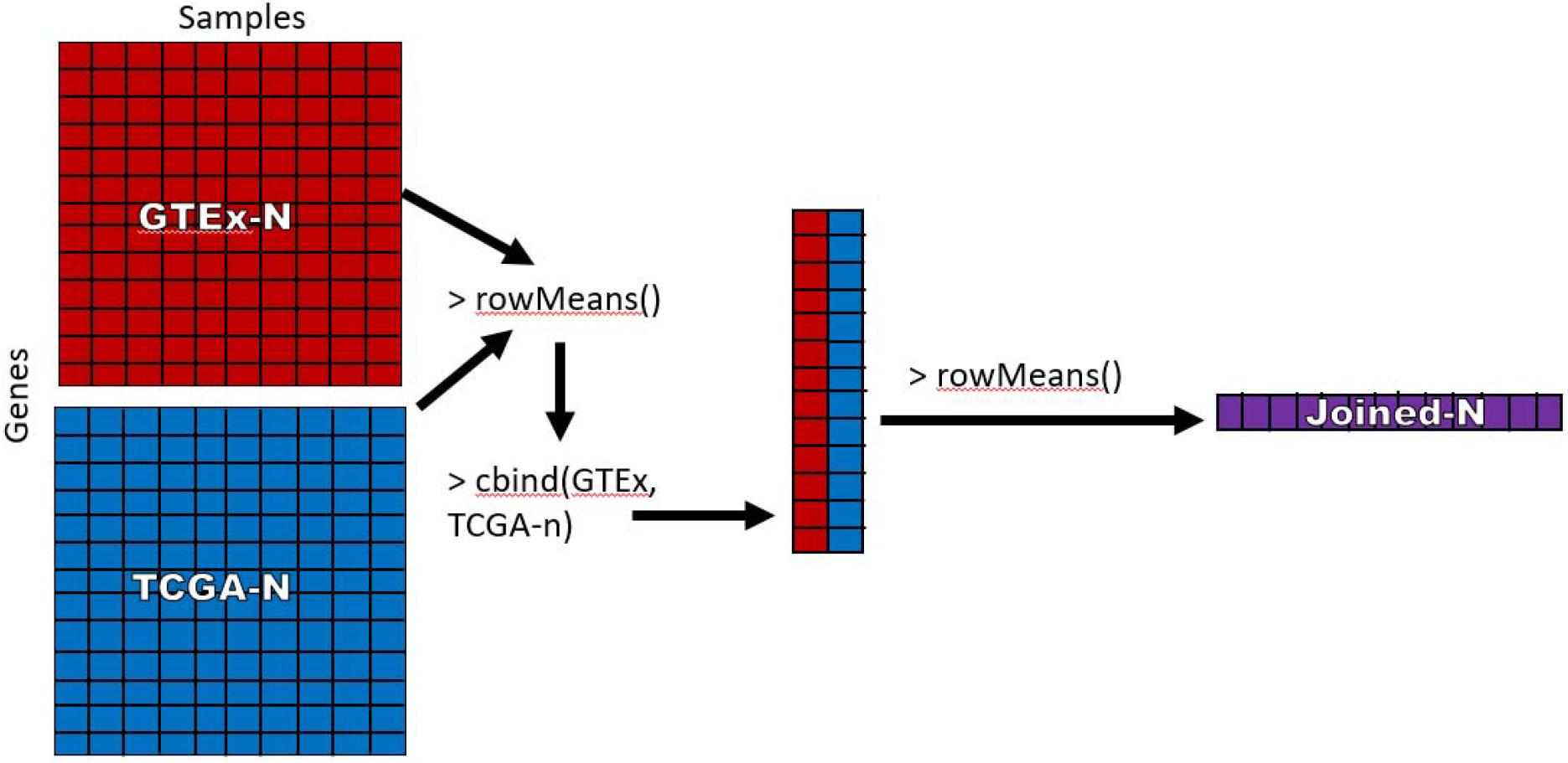
The generation of the Joined-N gene list using R code

**Figure S7:**
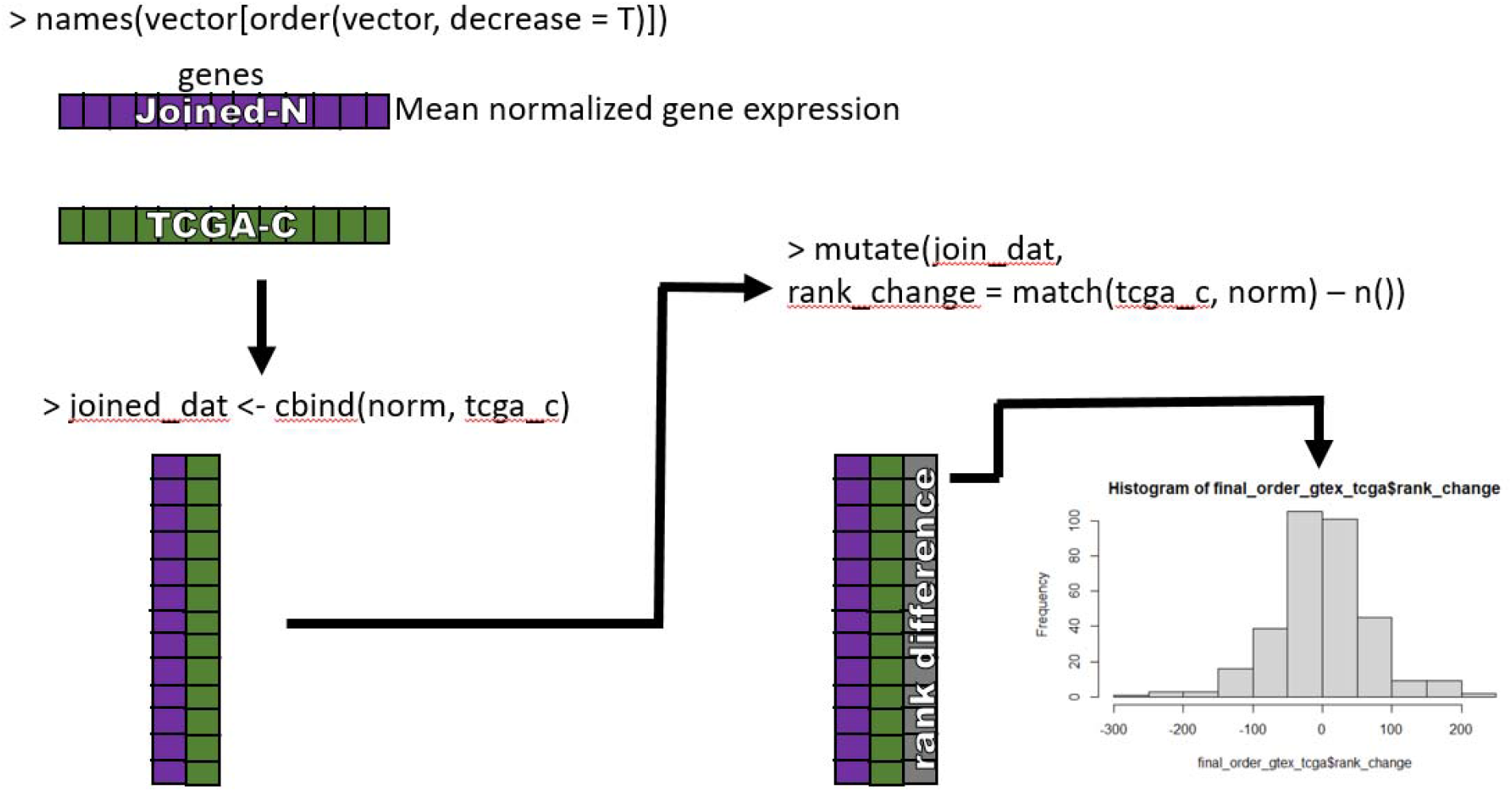
Generation of normal to cancer rank change using R code

## References

[1] C. Frantz, K.M. Stewart, V.M. Weaver, The extracellular matrix at a glance, J Cell Sci 123(Pt 24) (2010) 4195–200, doi: 10.1242/jcs.023820.

[2] A. Naba, K.R. Clauser, H. Ding, C.A. Whittaker, S.A. Carr, R.O. Hynes, The extracellular matrix: Tools and insights for the “omics” era, Matrix biology : journal of the International Society for Matrix Biology 49 (2016) 10–24, doi: 10.1016/j.matbio.2015.06.003.

[3] J.F. Bateman, R.P. Boot-Handford, S.R. Lamande, Genetic diseases of connective tissues: cellular and extracellular effects of ECM mutations, Nat Rev Genet 10(3) (2009) 173–83, doi: 10.1038/nrg2520.

[4] B.L. Loeys, J. Chen, E.R. Neptune, D.P. Judge, M. Podowski, T. Holm, J. Meyers, C.C. Leitch, N. Katsanis, N. Sharifi, F.L. Xu, L.A. Myers, P.J. Spevak, D.E. Cameron, J. De Backer, J. Hellemans, Y. Chen, E.C. Davis, C.L. Webb, W. Kress, P. Coucke, D.B. Rifkin, A.M. De Paepe, H.C. Dietz, A syndrome of altered cardiovascular, craniofacial, neurocognitive and skeletal development caused by mutations in TGFBR1 or TGFBR2, Nat Genet 37(3) (2005) 275–81, doi: 10.1038/ng1511.

[5] K.A. Piez, History of extracellular matrix: a personal view, Matrix biology : journal of the International Society for Matrix Biology 16(3) (1997) 85–92, doi: 10.1016/s0945-053x(97)90037-8.

[6] R.O. Hynes, A. Naba, Overview of the matrisome--an inventory of extracellular matrix constituents and functions, Cold Spring Harbor perspectives in biology 4(1) (2012) a004903, doi: 10.1101/cshperspect.a004903.

[7] A. Naba, K.R. Clauser, S. Hoersch, H. Liu, S.A. Carr, R.O. Hynes, The matrisome: in silico definition and in vivo characterization by proteomics of normal and tumor extracellular matrices, Molecular & cellular proteomics : MCP 11(4) (2012) M111 014647, doi: 10.1074/mcp.M111.014647.

[8] D. Wang, B. Eraslan, T. Wieland, B. Hallstrom, T. Hopf, D.P. Zolg, J. Zecha, A. Asplund, L.H. Li, C. Meng, M. Frejno, T. Schmidt, K. Schnatbaum, M. Wilhelm, F. Ponten, M. Uhlen, J. Gagneur, H. Hahne, B. Kuster, A deep proteome and transcriptome abundance atlas of 29 healthy human tissues, Mol Syst Biol 15(2) (2019) e8503, doi: 10.15252/msb.20188503.

[9] L. Jiang, M. Wang, S. Lin, R. Jian, X. Li, J. Chan, G. Dong, H. Fang, A.E. Robinson, G.T. Consortium, M.P. Snyder, A Quantitative Proteome Map of the Human Body, Cell 183(1) (2020) 269–283 e19, doi: 10.1016/j.cell.2020.08.036.

[10] A. Banazadeh, L. Veillon, K.M. Wooding, M. Zabet-Moghaddam, Y. Mechref, Recent advances in mass spectrometric analysis of glycoproteins, Electrophoresis 38(1) (2017) 162–189, doi: 10.1002/elps.201600357.

[11] T.O. Nieuwenhuis, S.Y. Yang, R.X. Verma, V. Pillalamarri, D.E. Arking, A.Z. Rosenberg, M.N. McCall, M.K. Halushka, Consistent RNA sequencing contamination in GTEx and other data sets, Nature communications 11(1) (2020) 1933, doi: 10.1038/s41467-020-15821-9.

[12] M.N. McCall, P.B. Illei, M.K. Halushka, Complex Sources of Variation in Tissue Expression Data: Analysis of the GTEx Lung Transcriptome, American journal of human genetics 99(3) (2016) 624–35, doi: 10.1016/j.ajhg.2016.07.007.

[13] M. Soumillon, A. Necsulea, M. Weier, D. Brawand, X. Zhang, H. Gu, P. Barthes, M. Kokkinaki, S. Nef, A. Gnirke, M. Dym, B. de Massy, T.S. Mikkelsen, H. Kaessmann, Cellular source and mechanisms of high transcriptome complexity in the mammalian testis, Cell reports 3(6) (2013) 2179–90, doi: 10.1016/j.celrep.2013.05.031.

[14] J.J. Maleszewski, C.K. Lai, J.P. Veinot, Anatomic Considerations and Examination of Cardiovascular Specimens (Excluding Devices), in: L.M. Buja, J. Butany (Eds.), Cardiovascular Pathology, Elsevier2015.

[15] M.J. Divo, C.H. Martinez, D.M. Mannino, Ageing and the epidemiology of multimorbidity, Eur Respir J 44(4) (2014) 1055–68, doi: 10.1183/09031936.00059814.

[16] M.I. Christodoulou, M. Avgeris, I. Kokkinopoulou, E. Maratou, P. Mitrou, C.K. Kontos, E. Pappas, E. Boutati, A. Scorilas, E.G. Fragoulis, Blood-based analysis of type-2 diabetes mellitus susceptibility genes identifies specific transcript variants with deregulated expression and association with disease risk, Scientific reports 9(1) (2019) 1512, doi: 10.1038/s41598-018-37856-1.

[17] G.R. Hunter, B.A. Gower, B.L. Kane, Age Related Shift in Visceral Fat, Int J Body Compos Res 8(3) (2010) 103–108, doi.

[18] M. Gershoni, S. Pietrokovski, The landscape of sex-differential transcriptome and its consequent selection in human adults, BMC Biol 15(1) (2017) 7, doi: 10.1186/s12915-017-0352-z.

[19] K. Karastergiou, S.R. Smith, A.S. Greenberg, S.K. Fried, Sex differences in human adipose tissues - the biology of pear shape, Biol Sex Differ 3(1) (2012) 13, doi: 10.1186/2042-6410-3-13.

[20] H. Dao, Jr., R.A. Kazin, Gender differences in skin: a review of the literature, Gend Med 4(4) (2007) 308–28, doi: 10.1016/s1550-8579(07)80061-1.

[21] S. Shuster, M.M. Black, E. McVitie, The influence of age and sex on skin thickness, skin collagen and density, Br J Dermatol 93(6) (1975) 639–43, doi: 10.1111/j.1365-2133.1975.tb05113.x.

[22] B. Mittal, Subcutaneous adipose tissue & visceral adipose tissue, Indian J Med Res 149(5) (2019) 571–573, doi: 10.4103/ijmr.IJMR_1910_18.

[23] N. Cancer Genome Atlas Research, J.N. Weinstein, E.A. Collisson, G.B. Mills, K.R. Shaw, B.A. Ozenberger, K. Ellrott, I. Shmulevich, C. Sander, J.M. Stuart, The Cancer Genome Atlas Pan-Cancer analysis project, Nat Genet 45(10) (2013) 1113–20, doi: 10.1038/ng.2764.

[24] S. Fujishima, T. Shiomi, S. Yamashita, Y. Yogo, Y. Nakano, T. Inoue, M. Nakamura, S. Tasaka, N. Hasegawa, N. Aikawa, A. Ishizaka, Y. Okada, Production and activation of matrix metalloproteinase 7 (matrilysin 1) in the lungs of patients with idiopathic pulmonary fibrosis, Archives of pathology & laboratory medicine 134(8) (2010) 1136–42, doi: 10.1043/2009-0144-OA.1.

[25] T.S. Adams, J.C. Schupp, S. Poli, E.A. Ayaub, N. Neumark, F. Ahangari, S.G. Chu, B.A. Raby, G. DeIuliis, M. Januszyk, Q. Duan, H.A. Arnett, A. Siddiqui, G.R. Washko, R. Homer, X. Yan, I.O. Rosas, N. Kaminski, Single-cell RNA-seq reveals ectopic and aberrant lung-resident cell populations in idiopathic pulmonary fibrosis, Sci Adv 6(28) (2020) eaba1983, doi: 10.1126/sciadv.aba1983.

[26] P. Sivakumar, J.R. Thompson, R. Ammar, M. Porteous, C. McCoubrey, E. Cantu, 3rd, K. Ravi, Y. Zhang, Y. Luo, D. Streltsov, M.F. Beers, G. Jarai, J.D. Christie, RNA sequencing of transplant-stage idiopathic pulmonary fibrosis lung reveals unique pathway regulation, ERJ Open Res 5(3) (2019) doi: 10.1183/23120541.00117-2019.

[27] M. Mele, P.G. Ferreira, F. Reverter, D.S. DeLuca, J. Monlong, M. Sammeth, T.R. Young, J.M. Goldmann, D.D. Pervouchine, T.J. Sullivan, R. Johnson, A.V. Segre, S. Djebali, A. Niarchou, G.T. Consortium, F.A. Wright, T. Lappalainen, M. Calvo, G. Getz, E.T. Dermitzakis, K.G. Ardlie, R. Guigo, Human genomics. The human transcriptome across tissues and individuals, Science 348(6235) (2015) 660–5, doi: 10.1126/science.aaa0355.

[28] H. Wulf-Johansson, S. Lock Johansson, A. Schlosser, A. Trommelholt Holm, L.M. Rasmussen, H. Mickley, A.C. Diederichsen, H. Munkholm, T.S. Poulsen, I. Tornoe, V. Nielsen, N. Marcussen, J. Vestbo, S.G. Saekmose, U. Holmskov, G.L. Sorensen, Localization of microfibrillar-associated protein 4 (MFAP4) in human tissues: clinical evaluation of serum MFAP4 and its association with various cardiovascular conditions, PLoS One 8(12) (2013) e82243, doi: 10.1371/journal.pone.0082243.

[29] S.G. Saekmose, B. Mossner, P.B. Christensen, K. Lindvig, A. Schlosser, R. Holst, T. Barington, U. Holmskov, G.L. Sorensen, Microfibrillar-Associated Protein 4: A Potential Biomarker for Screening for Liver Fibrosis in a Mixed Patient Cohort, PLoS One 10(10) (2015) e0140418, doi: 10.1371/journal.pone.0140418.

[30] S. Wada, Y. Yasunaga, K. Oka, N. Dan, E. Tanaka, K. Morita, E. Masuda, K. Yanagawa, H. Matsumoto, S. Yoshioka, M. Tsujie, Y. Inui, S. Kawata, Submucosal fat accumulation in human colorectal tissue and its association with abdominal obesity and insulin resistance, United European Gastroenterol J 6(7) (2018) 1065–1073, doi: 10.1177/2050640618766926.

[31] H. Mesa, S. Drawz, R. Dykoski, J.C. Manivel, Morphometric measurement of submucosal thickness in areas of fat deposition in the terminal ileum and colonic sections, with correlation with body mass index, weight and age: a male autopsy study, Histopathology 67(4) (2015) 457–63, doi: 10.1111/his.12683.

[32] J. Collazos, V. Asensi, G. Martin, A.H. Montes, T. Suarez-Zarracina, E. Valle-Garay, The effect of gender and genetic polymorphisms on matrix metalloprotease (MMP) and tissue inhibitor (TIMP) plasma levels in different infectious and non-infectious conditions, Clinical and experimental immunology 182(2) (2015) 213–9, doi: 10.1111/cei.12686.

[33] A. Samnegard, A. Silveira, P. Lundman, S. Boquist, J. Odeberg, J. Hulthe, W. McPheat, P. Tornvall, L. Bergstrand, C.G. Ericsson, A. Hamsten, P. Eriksson, Serum matrix metalloproteinase-3 concentration is influenced by MMP-3 -1612 5A/6A promoter genotype and associated with myocardial infarction, Journal of internal medicine 258(5) (2005) 411–9, doi: 10.1111/j.1365-2796.2005.01561.x.

[34] S.J. Fitzgerald, A.V. Janorkar, A. Barnes, R.O. Maranon, A new approach to study the sex differences in adipose tissue, J Biomed Sci 25(1) (2018) 89, doi: 10.1186/s12929-018-0488-3.

[35] K.B. Chapman, M.J. Prendes, H. Sternberg, J.L. Kidd, W.D. Funk, J. Wagner, M.D. West, COL10A1 expression is elevated in diverse solid tumor types and is associated with tumor vasculature, Future Oncol 8(8) (2012) 1031–40, doi: 10.2217/fon.12.79.

[36] D. Jia, Z. Liu, N. Deng, T.Z. Tan, R.Y. Huang, B. Taylor-Harding, D.J. Cheon, K. Lawrenson, W.R. Wiedemeyer, A.E. Walts, B.Y. Karlan, S. Orsulic, A COL11A1-correlated pan-cancer gene signature of activated fibroblasts for the prioritization of therapeutic targets, Cancer letters 382(2) (2016) 203–214, doi: 10.1016/j.canlet.2016.09.001.

[37] G.G. Seniski, A.A. Camargo, D.F. Ierardi, E.A. Ramos, M. Grochoski, E.S. Ribeiro, I.J. Cavalli, F.O. Pedrosa, E.M. de Souza, S.M. Zanata, F.F. Costa, G. Klassen, ADAM33 gene silencing by promoter hypermethylation as a molecular marker in breast invasive lobular carcinoma, BMC cancer 9 (2009) 80, doi: 10.1186/1471-2407-9-80.

[38] G.C. Manica, C.F. Ribeiro, M.A. Oliveira, I.T. Pereira, A. Chequin, E.A. Ramos, L.M. Klassen, A.P. Sebastiao, L.M. Alvarenga, S.M. Zanata, L. Noronha, I. Rabinovich, F.F. Costa, E.M. Souza, G. Klassen, Down regulation of ADAM33 as a Predictive Biomarker of Aggressive Breast Cancer, Scientific reports 7 (2017) 44414, doi: 10.1038/srep44414.

[39] M. Yamatoji, A. Kasamatsu, Y. Kouzu, H. Koike, Y. Sakamoto, K. Ogawara, M. Shiiba, H. Tanzawa, K. Uzawa, Dermatopontin: a potential predictor for metastasis of human oral cancer, Int J Cancer 130(12) (2012) 2903–11, doi: 10.1002/ijc.26328.

[40] A.M. Macgregor, C.G. Eberhart, M. Fraig, J. Lu, M.K. Halushka, Tissue inhibitor of matrix metalloproteinase-3 levels in the extracellular matrix of lung, kidney, and eye increase with age, J Histochem Cytochem 57(3) (2009) 207–13, doi: 10.1369/jhc.2008.952531.

[41] B. Schwanhausser, D. Busse, N. Li, G. Dittmar, J. Schuchhardt, J. Wolf, W. Chen, M. Selbach, Global quantification of mammalian gene expression control, Nature 473(7347) (2011) 337–42, doi: 10.1038/nature10098.

[42] M. Uhlen, L. Fagerberg, B.M. Hallstrom, C. Lindskog, P. Oksvold, A. Mardinoglu, A. Sivertsson, C. Kampf, E. Sjostedt, A. Asplund, I. Olsson, K. Edlund, E. Lundberg, S. Navani, C.A. Szigyarto, J. Odeberg, D. Djureinovic, J.O. Takanen, S. Hober, T. Alm, P.H. Edqvist, H. Berling, H. Tegel, J. Mulder, J. Rockberg, P. Nilsson, J.M. Schwenk, M. Hamsten, K. von Feilitzen, M. Forsberg, L. Persson, F. Johansson, M. Zwahlen, G. von Heijne, J. Nielsen, F. Ponten, Proteomics. Tissue-based map of the human proteome, Science 347(6220) (2015) 1260419, doi: 10.1126/science.1260419.

[43] C.A. Schneider, W.S. Rasband, K.W. Eliceiri, NIH Image to ImageJ: 25 years of image analysis, Nature methods 9(7) (2012) 671–5, doi.

[44] M.E. Ritchie, B. Phipson, D. Wu, Y. Hu, C.W. Law, W. Shi, G.K. Smyth, limma powers differential expression analyses for RNA-sequencing and microarray studies, Nucleic Acids Res 43(7) (2015) e47, doi: 10.1093/nar/gkv007.

